# Chromosome segregation dynamics during the cell cycle of *Staphylococcus aureus*

**DOI:** 10.1101/2025.02.18.638847

**Authors:** Adrian Izquierdo-Martinez, Simon Schäper, António D. Brito, Qin Liao, Coralie Tesseur, Moritz Sorg, Daniela S. Botinas, Xindan Wang, Mariana G. Pinho

## Abstract

Research on chromosome organization and cell cycle progression in spherical bacteria, particularly *Staphylococcus aureus*, remains limited and fragmented. In this study, we established a working model to investigate chromosome dynamics in *S. aureus* using a Fluorescent Repressor-Operator System (FROS), which enabled precise localization of specific chromosomal loci. This approach revealed that the *S. aureus* cell cycle and chromosome replication cycle are not coupled, with cells exhibiting two segregated origins of replication at the start of the cell cycle. The chromosome has a specific origin-terminus-origin conformation, with origins localizing near the membrane, towards the tip of each hemisphere, or the “cell poles”. We further used this system to assess the role of various proteins with a role in *S. aureus* chromosome biology, focusing on the ParB-*parS* and SMC-ScpAB systems. Our results demonstrate that ParB binds five *parS* chromosomal sequences and the resulting complexes influence chromosome conformation, but play a minor role in chromosome compaction and segregation. In contrast, the SMC-ScpAB complex plays a key role in *S. aureus* chromosome biology, contributing to chromosome compaction, segregation and spatial organization. Additionally, we systematically assessed and compared the impact of proteins linking chromosome segregation to cell division—Noc, FtsK, SpoIIIE and XerC—on origin and terminus number and positioning. This work provides a comprehensive study of the factors governing chromosome dynamics and organization in *S. aureus*, contributing to our knowledge on chromosome biology of spherical bacteria.

## Introduction

Bacteria exhibit an exquisite spatiotemporal organization of cellular components. From generating their own shape, to dividing with precision and placing structures and organelles in specific locations, these microorganisms coordinate their cellular activities and machinerýs localization with remarkable accuracy. One key example is chromosome dynamics, which includes the spatial organization of the chromosome, its replication and proper segregation into daughter cells, all of which are essential for the faithful transmission of genetic information to the next generation (*1*, *2*).

Studies on multiple bacterial model organisms have revealed specific arrangements for the chromosome during the processes of replication and segregation (Fig. S1). Some organisms like *Caulobacter crescentus* (*3*), *Myxococcus xanthus* (*4*), or the chromosome 1 of *Vibrio cholerae* (*5*) display an organization known as linear (Ori-Ter) chromosome arrangement in which the replication origin (Ori) is in a polar (sub-polar in *M. xanthus*) region and the replication terminus (Ter) sits on the opposite pole in newborn cells. After the initiation of chromosome replication, one of the newly replicated Ori is segregated to the opposite cell pole, and the Ter region moves to the cell center, near the division site, creating an Ori-Ter-Ori arrangement. Such dynamics ensures that both daughter cells inherit a copy of the chromosome which is in the same orientation as in the mother cell.

Newborn cells of the model organism *Bacillus subtilis,* in slow-growing conditions, also present an Ori-Ter-Ori orientation of partially replicated chromosomes. However, after their complete replication, *B. subtilis* chromosomes adopt a Left-Ori-Right configuration, with the Ori and Ter regions of each chromosome in the ¼ and ¾ positions of the cell (*6*, *7*). As chromosome replication re-starts before the end of the cell cycle, the origins segregate to the poles and the septum regions (Fig. S1). The actinobacterium *Corynebacterium glutamicum* has a similar cycle, but starting with two completely replicated chromosomes, and the origins remain in the polar positions during the entire cell cycle (*8*). On the other hand, *Escherichia coli* in slow-growing conditions adopts a Left-Ori-Right organization, with the origin in the cell center in newborn cells (*9–12*). After the initiation of replication, the origins segregate to the ¼ and ¾ positions, where they remain until replication is finished and the cells divide (Fig. S1). Although phylogenetically distant to *E. coli*, a similar chromosomal arrangement can be found in both the ovococcoid firmicute *Streptococcus pneumoniae* (*13*) and the actinobacterium *Mycobacterium smegmatis* (*14*). Besides the diversity of chromosome arrangements, an additional factor that adds complexity to the process of chromosome segregation is the presence of multifork replication, that is, the initiation of new replication rounds in chromosomes that are still being replicated, which increases the number of origins per cell. This has been observed in some organisms, such as *B. subtilis* (*15*) and *E. coli* (*16*) in fast growth conditions, and in the slow-growing *M. smegmatis* (*17*).

Chromosome organization is highly dependent on the correct segregation of the origin regions, which is mediated by multiple factors that vary among bacterial species. In *B. subtilis* (*18*), *C. crescentus* (*19*), *V. cholerae* (*20*) or *M. xanthus* (*21*), among others, ParABS systems play a direct role in origin segregation. In brief, the ParB component of the system is a CTPase that interacts with itself and with *parS* sequences generally located near the origin, forming a kinetochore-like structure that can be mobilized through cyclic dimerization and monomerization of the ParA component, an ATPase whose dimers interact both with ParB and the DNA (*22*). There is evidence that these systems might be present in the majority of bacterial species (*23*). A second function of ParB is to load SMC-ScpAB condensin-like complexes onto chromosomal *parS* sites (*24–26*). These complexes contribute to the overall compaction of the nucleoid and to the segregation of sister chromosomes as they are being replicated (*27–30*). SMC molecules dimerize spontaneously, forming a V-shaped structure, with long coiled-coil arms connecting two complete ATPase domains (or heads), formed by the C- and N-terminus of each monomer, to a central hinge. ScpA interacts with SMC by bridging the two heads of the dimer, forming a ring-like structure that is able to entrap DNA, while ScpB dimers associate to ScpA and are required for the loading of SMC-ScpAB complexes onto the chromosome (*31*, *32*). The current model proposes that ParB-*parS* nucleocomplexes load the SMC complexes onto the chromosome, which then travel from the origin region to the terminus, juxtaposing and aligning the two chromosome arms and therefore contributing to the overall spatial organization and compaction of the chromosome (*26*, *29*, *30*, *33–35*). Depending on the bacterial species, the role of SMC-ScpAB or ParABS systems can vary from being essential for survival (particularly in fast growing conditions) to their absence causing only mild phenotypes (*19*, *21*, *33*, *34*, *36–43*). Besides specific proteins with a role in chromosome segregation, the physico-chemical properties of the chromosome are proposed to contribute to the spontaneous unmixing of sister chromatids (*44*). In turn, chromosome organization plays a role in the regulation of other cellular processes, like cell division. In many organisms, nucleoid-associated proteins prevent the progression of septum-formation, coordinating both processes and preventing guillotining of the chromosome (*45–49*).

Most knowledge on chromosome segregation comes from a selected group of rod-shaped model organisms. The geometry of the bacterium is a key player in its spatial organization, as the presence of topologically different regions allows to determine separate spaces in the cell (*50*, *51*). For example, proteins sensing curvature are involved in chromosome positioning (*52*), division site placement (*53*, *54*) and cell morphogenesis (*55–57*). Therefore, the existence of spherical bacteria which, during part of the cell cycle, appear to have a constant curvature in every direction of their surfaces, raises questions about how their chromosomes are spatially organized and how this organization is maintained.

The model organism used in this work, *Staphylococcus aureus*, is a firmicute with a nearly spherical morphology. *S. aureus* is a human opportunistic pathogen that generally inhabits the skin as a commensal but can cause a variety of infections. Moreover, many *S. aureus* strains have acquired resistance to beta-lactam antibiotics, and some pathogenic strains are resistant to most antibiotic classes (*58*). Its widespread presence in the human population, combined with its drug resistance, makes *S. aureus* a significant threat to human health and a major cause of death from antibiotic-resistant infections (*59*). Besides its clinical relevance, *S. aureus* is an important model organism in the bacterial cell biology field, as it is one of the few intensively studied coccoid organisms. Its cell cycle is divided into three stages (*60*, *61*): it begins with Phase 1 (P1), characterized by a nearly spherical newborn cell. As the cell starts to build a septum, it transitions into Phase 2 (P2), during which the DNA must be segregated into each of the developing compartments or hemispheres. Previous data suggest that septum formation is determined by the orientation of the segregated chromosomes, linking nucleoid spatial organization with cell division (*47*, *62*). Once the septum is completed, the cell enters Phase 3 (P3), characterized by two compartmentalized cytoplasmic spaces. At the end of P3 the septum splits rapidly, giving rise to two P1 cells.

Dynamics of chromosome segregation in *S. aureus* is currently understudied, with most information available deriving from studies of ParB localization as a proxy for Ori localization (*47*, *63*, *64*). In the present study, we localized specific chromosomal loci in *S. aureus* and showed that newborn cells generally have a partially replicated chromosome with two segregated origins. We show that the origin has a preferential localization pattern towards the tip of each hemisphere, henceforth referred to as “cell poles”, while the terminus is restricted to the cell center, resulting in an Ori-Ter-Ori chromosomal organization. Furthermore, we systematically analyzed the role of proteins known to influence chromosome segregation/dynamics, offering a comprehensive understanding of the factors contributing to chromosome biology of *S. aureus*.

## Results

### *S. aureus* cell cycle and chromosome replication cycle are not coupled

The number, orientation and dynamics of the *S. aureus* chromosome have not been comprehensively investigated. Localization studies of fluorescent derivatives of ParB (also known as Spo0J) show that cells generally have two to four origins (*47*, *62*, *63*). One limitation of these studies is that, in our hands, fluorescently tagged versions of ParB produce dim and poorly condensed foci, not ideal for a precise analysis (*47*). Therefore, we adapted a fluorescent repressor operator system (FROS) initially developed for *E. coli* (*11*) for the visualization of specific chromosomal loci in *S. aureus*. This system makes use of fluorescent derivatives of the Lac repressor (LacI) and the Tet repressor (TetR) that bind to, and allow visualization of, arrays containing 48 copies of *lacO* or *tetO* operator sequences, respectively, which are introduced at specific locations on the bacterial chromosome. For this purpose, we first introduced the *lacO* and *tetO* arrays at chromosomal loci near the Ori or Ter regions. We then introduced the *lacI* and *tetR* genes fused to sequences encoding the fluorescent proteins eCFP and eYFP (*11*), respectively, under the control of a cadmium-inducible promoter (*65*) in the *spa* locus of *S. aureus* JE2 chromosome (*66*), generating strains JE2_Ori_CFP_Ter_YFP and JE2_Ori_YFP_Ter_CFP which allowed us to simultaneously visualize two different loci in each strain. We tested two different cadmium induction times and two combinations of operator arrays, and observed the formation of fluorescent foci corresponding to Ori and Ter localization in the cells (Fig. S2). Cells usually had two to four origins, similar to what had been reported using a ParB fluorescent derivative in *S. aureus* (*47*, *62*, *63*). However, we noticed that the average number of foci per cell varied slightly depending on the duration of cadmium induction and the type of operator sequence array (*tetO*/*lacO*) used to label each region (Fig. S2B). Such variability could compromise quantitative studies comparing different strains. Fortuitously, we noticed that a fusion of mNeonGreen (*67*) to TetR (TetR-mNG, strain JE2_FROS^ori^) was sufficiently expressed in the absence of inducer to allow clear foci formation. Thus, we used this constitutive single-locus FROS system (Fig. 1A) when labelling only one chromosomal position was sufficient, eliminating variations due to induction time and/or to cellular responses to the presence of cadmium.

**Figure 1.**
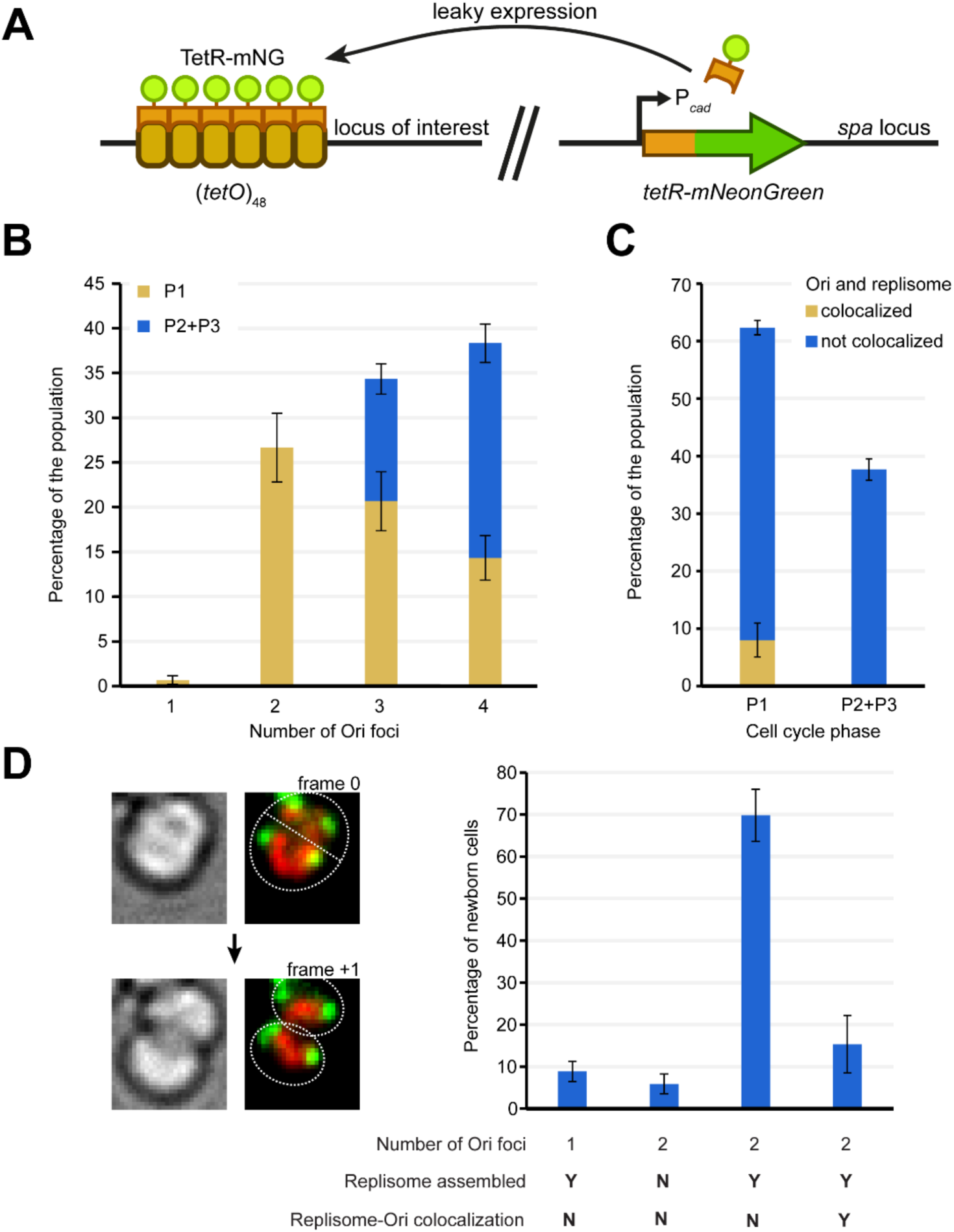
Single-Locus FROS reveals number of origins across cell cycle stages. **A** Schematic of the single-locus FROS system used in this study. A sequence encoding the TetR repressor fused to the mNeonGreen (mNG) fluorescent protein, under the control of a cadmium-inducible promoter with leaky expression, was integrated in the *spa* locus (right). TetR-mNG binds to an array of 48 *tetO* operator sequences introduced at the locus of interest (left). **B** Bar chart representing the relative frequency of cells of the JE2_FROS^ori^_DnaN-Halo strain with 1-4 Ori foci in P1 (yellow, no septum) or P2/P3 (blue, incomplete and complete septum) cell cycle stages. Data from three biological replicates (n=100 each), error bars indicate standard deviation. **C** Bar chart showing the relative frequency of P1 and P2/P3 cells of the JE2_FROS^ori^_DnaN-Halo strain with Ori and replisome (visualized using DnaN as a proxy) either colocalizing (yellow) or not (blue). Data from three biological replicates (n=100 each), error bars indicate standard deviation. **D** Classification of newborn cells by Ori number and Ori/replisome colocalization. Left, brightfield and fluorescence microscopy images of a representative cell from JE2_FROS^ori^_DnaN-Halo strain that underwent division between frame 0 and frame +1, showing two newborn cells with two Ori each in the latter frame (3 min interval between frames). JF549-labelled DnaN-Halo signal in red and TetR-mNG (Ori) signal in green. Right, bar chart showing the relative frequency of each class of newborn cells, categorized by Ori number, replisome assembly, and Ori/replisome colocalization. Data from three biological replicates (n= 36, 36, 30), error bars indicate standard deviation.

In addition to the FROS system to localize different chromosomal loci, we constructed a HaloTag fusion (*68*) to DnaN, a component of the replisome whose localization has been used as a proxy for the replisome in studies of other organisms (*42*, *69*). These tools allowed us to follow cellular localization of chromosome loci and of the replisome, enabling us to characterize the chromosome replication cycle of *S. aureus*.

In a first approach, we classified *S. aureus* cells according to the number of origin foci and correlated that information with the cell cycle stage. When quantifying the number of origins in a spherical cell, two factors can result in an underestimation: (i) two foci separated by a distance smaller than the resolution limit appear as a single focus; (ii) two foci with similar coordinates in the *xy* plane (parallel to the microscope slide) but different coordinate in the *z* axis (perpendicular to the microscope slide) appear as a single focus in a microscopy image. To overcome the latter limitation, we imaged JE2_FROS^ori^_DnaN-Halo cells in three Z-planes followed by manual analysis of each plane to count all detected foci (Fig. 1B and S3). Using this approach, we rarely observed cells with a single Ori focus (<1%), with cells typically having two (∼27%), three (∼34%) or four (∼38%) Ori foci (Fig. 1B). As expected, the number of origins increases as the cell cycle progresses, with P2 and P3 cells having three or four Ori foci, indicating that four origins are the typical maximum.

We then asked when, during the cell cycle, is replication initiated. For that, we assessed colocalization of the origin and the replisome protein DnaN in strain JE2_FROS^ori^_DnaN-Halo. This colocalization was only observed in P1 cells, suggesting that re-initiation of chromosome replication happens during that stage (Fig. 1C). Furthermore, the replisome was assembled, i.e. formed one or more foci (as opposed to having a diffuse cytoplasmic signal) in >95% of the cells (Fig. S3), indicating that DNA replication occurs during almost the entire cell cycle.

To directly show, in time-lapse movies, that cells start their cell cycle with two segregated origins of replication, JE2_FROS^ori^_DnaN-Halo cells were labelled with red fluorescent dye JF549-HTL and imaged every 3 minutes. By analyzing the frame immediately after the splitting of the mother cell in two daughter cells, we showed that more than 90% of newborn cells had two Ori foci (Fig. 1D). Additionally, around 70% of cells had an assembled replisome that did not colocalize with the origins, indicating that cells are typically born with a partially replicated chromosome (Fig. 1D). We have also observed newborn cells (∼20%) containing two origins either with a diffuse DnaN signal (no active replisome, indicating that the replication round is finished) or with DnaN colocalizing with the Ori foci (replisome initiating a next round of replication), indicating that in these cases cells are born with two completely replicated chromosomes (Fig. 1D). When following the cell cycle in time-lapse experiments, we could also identify cells with two Ori that underwent the completion of one round of replication and initiated a new round, followed by origin segregation, resulting in cells with four Ori foci (Fig. S4). These observations further support that, during its cell cycle, *S. aureus* generally progresses from a partially replicated chromosome to two partially replicated chromosomes.

Collectively, the data show that the *S. aureus* cell cycle (from one cell division to the next) and chromosome replication cycle (a complete round of chromosome replication) are not coupled: during a single cell cycle, most cells complete one replication cycle and initiate another, resulting in cells with four origins and two partially replicated chromosomes that will be distributed to the two daughter cells. This chromosome replication cycle resembles what has been described for *B. subtilis* under slow-growing conditions (*6*).

### *S. aureus* chromosome adopts a linear Ori-Ter-Ori conformation

Bacterial species generally have specific spatial arrangements of their chromosomes, particularly regarding the position of the Ori and Ter regions (Fig. S1). To study chromosome organization in *S. aureus*, we used the FROS system to label not only the Ori and Ter, but also the left and right arms of the chromosome (Fig. 2A, B). To systematically analyze the localization of labelled chromosome loci in thousands of cells, we developed a pipeline using the e-Hooke software version 1.1 (*70*) for cell segmentation, and TrackMate (*71*) for foci detection. This pipeline uses a maximum intensity projection of the TetR-mNG foci signal obtained from three Z-planes. Foci detection data is then used to generate average maps of the locations of different chromosomal regions within the cell (Fig. 2C). To generate these maps, cells were first aligned to the slightly longer axis, which is perpendicular to the future division plane. The relative position of each focus center in individual cells was recorded and mapped onto a model cell with median dimensions for length and width specific for each dataset. Heat maps were then created by calculating the average localization of foci, which correlates with the probability of a focus being found at each location. The resulting data showed that origins are typically positioned at the cell periphery, on opposite ends of the longer cell axis (the cell poles), termini are confined to the cell center, and the left and right arms occupy intermediate locations (Fig. 2C). These findings indicate that *S. aureus* cells adopt an Ori-Ter-Ori organization, with origins located in close proximity to the cell periphery.

**Figure 2.**
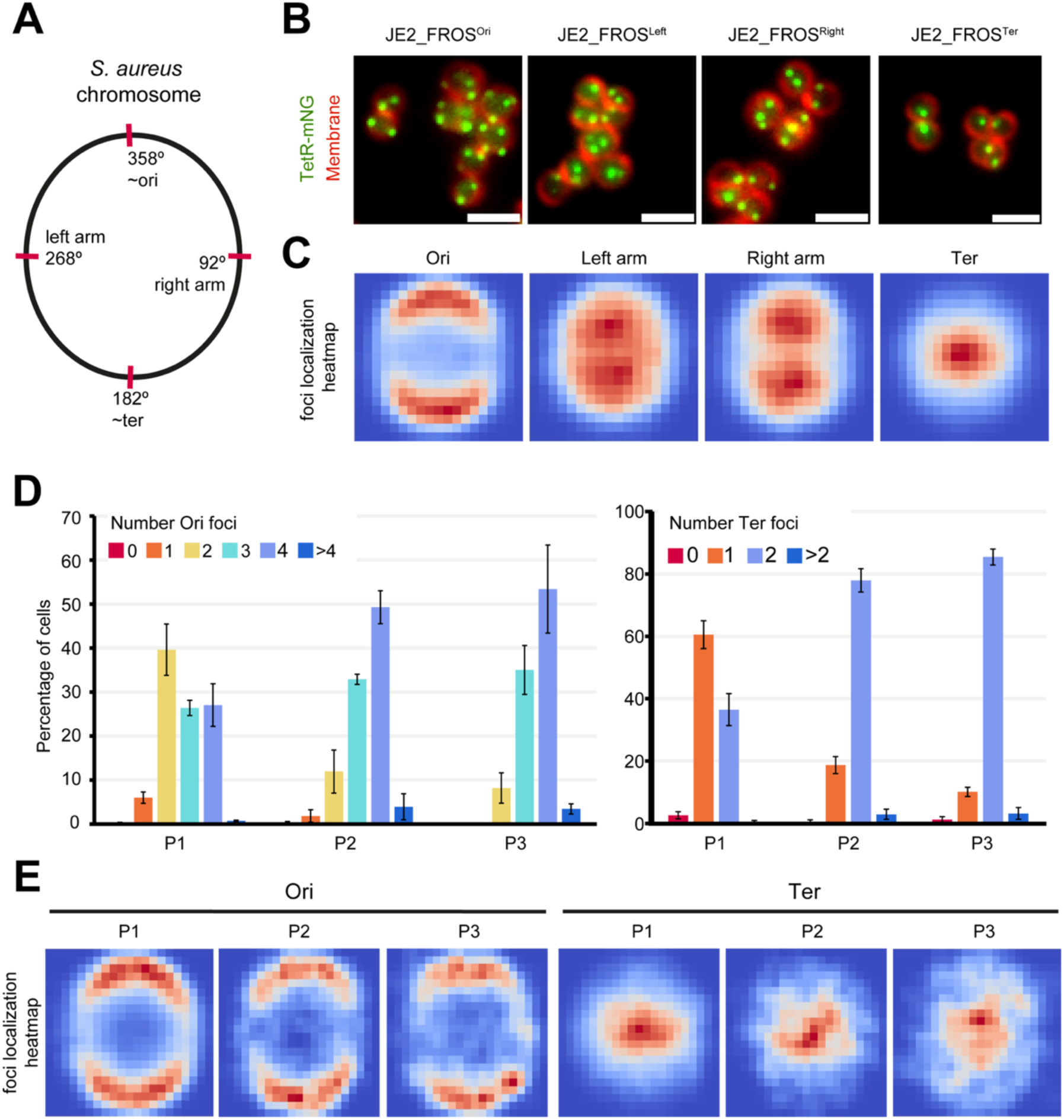
Localization of chromosomal regions during the cell cycle. **A** Schematic of the *S. aureus* chromosome showing the locations of (*tetO*)_48_ arrays used to localize the chromosomal origin, terminus, left arm and right arm. **B** Fluorescence microscopy images of cells with indicated chromosomal regions labeled by TetR-mNG (green) and membrane labeled with FM 4-64 dye (red). Scale bar: 2μm. **C** Heatmaps showing the average localization of detected fluorescent TetR-mNG spots in strains indicated in B. The color scale in each dataset ranges from red (maximum spot density) to dark blue (no spots detected), n>2500 cells per dataset. **D** Bar charts showing the distribution of the number of origin (left, strain JE2_FROS^Ori^) or terminus (right, strain JE2_FROS^Ter^) foci for cells in each cell phase of the cell cycle. Data from three biological replicates (n>600 each), error bars indicate standard deviation. **E** Heatmaps showing the average localization of detected fluorescence spots for cells of strains JE2_FROS^Ori^ (Ori) and JE2_FROS^Ter^ (Ter) in each phase of the cell cycle. Color scale as in panel C. From left to right n=3036, 1099, 436 (origin heatmaps); 1956, 610, 256 (terminus heatmaps).

The analyzed cells of the JE2_FROS^Ori^ and JE2_FROS^Ter^ strains were also manually classified according to their cell cycle phase, as described in (*60*), allowing us to quantify the number of origins and termini per cell at each cell cycle phase (Fig. 2D). Similarly to the data shown in Fig. 1B, we observed that P1 cells had two (∼40%), three (∼25%) or four (∼25%) origins, with the number of Ori per cell increasing as the cell cycle progresses (Fig. 2D). As for the termini, the majority of P1 cells had a single focus (∼60%), while most P2 and P3 cells had 2 foci (∼80%) (Fig. 2D). We also generated heatmaps of cells automatically classified according to their cell cycle phase (Fig. 2E), which showed that the Orís polar localization and the Teŕs central positioning were maintained throughout the cell cycle. These results confirm that *S. aureus* cells generally start the cell cycle with two origins, indicating that origin segregation usually occurs before septum synthesis is initiated. As the cell cycle progresses, a new round of chromosome replication produces cells with four origins and two termini.

### SMC complex, but not ParB, plays a major role in chromosome segregation in *S. aureus*

Having established the spatial organization and dynamics of the staphylococcal chromosome, we aimed to investigate the role of proteins reported to be involved in bacterial chromosome segregation and/or organization, specifically ParB and SMC. ParB is a key component of the ParABS system in various bacterial species (*22*). However, no ParA homolog has been identified in *S. aureus*, meaning that its ParABS system is incomplete, which makes the role of ParB in this bacterium particularly intriguing. It has been previously assumed that, similar to other ParABS systems, *S. aureus* ParB binds to Ori-proximal *parS* sequences, with one study predicting four such sequences in its genome (*23*). To experimentally determine the *parS* sites, we performed ChIP-Seq analysis using JE2_ParB-3xFLAG strain. We identified ParB enrichment at five loci, all near the Ori region (Fig. S5A). At each of these five locations, we found a *parS* motif, three of which (*parS*1, *parS*3 and *parS*5) coincide with those previously predicted (*23*) (Fig. S5B). Moreover, each ParB enrichment peak spanned 8-10 kb encompassing the *parS* sequence, indicating that, like in other organisms, *S. aureus* ParB nucleates around origin-proximal *parS* sequences and spreads to neighboring regions.

To understand the role of ParB and the SMC complex on the global organization of the chromosome, we used chromosome conformation capture (Hi-C) assay (*72*), a technique that involves crosslinking nearby DNA regions to capture the chromosome conformation by detecting the frequency of interaction between DNA loci across the whole genome. The *S. aureus* chromosome contact map for JE2 wild type strain (Fig. 3) displays a primary diagonal and a secondary diagonal. The primary diagonal has stronger signal, resulting from a high frequency of contacts between adjacent sequences in the chromosome, while the secondary diagonal has weaker signal, resulting from inter-arm, long distance DNA contacts. This interaction pattern supports that the chromosome has an Ori-Ter linear organization. Upon deletion of *parB,* the secondary diagonal disappears, consistent with previous findings (*73*), presumably because SMC is no longer loaded at specific *parS* loci, which is required for a constant set of inter-arm interactions. Furthermore, deletion or mutation of the five identified *parS* sites also abolished the secondary diagonal, confirming that both ParB and *parS* form a functional unit for chromosome organization (Fig. 3). Importantly, we constructed the strain JE2_Δ4*parS,* in which 4 *parS* sites were deleted or mutated and *parS*2 remained as the sole *parS* on the chromosome. This strain showed a more defined secondary diagonal compared to the wild type JE2, shifted towards the right arm, where *parS*2 is located (Fig. 3). Presumably, this occurs because SMC is loaded onto the chromosome from the single *parS*2 locus, resulting in a specific subset of inter-arm interactions.

**Figure 3.**
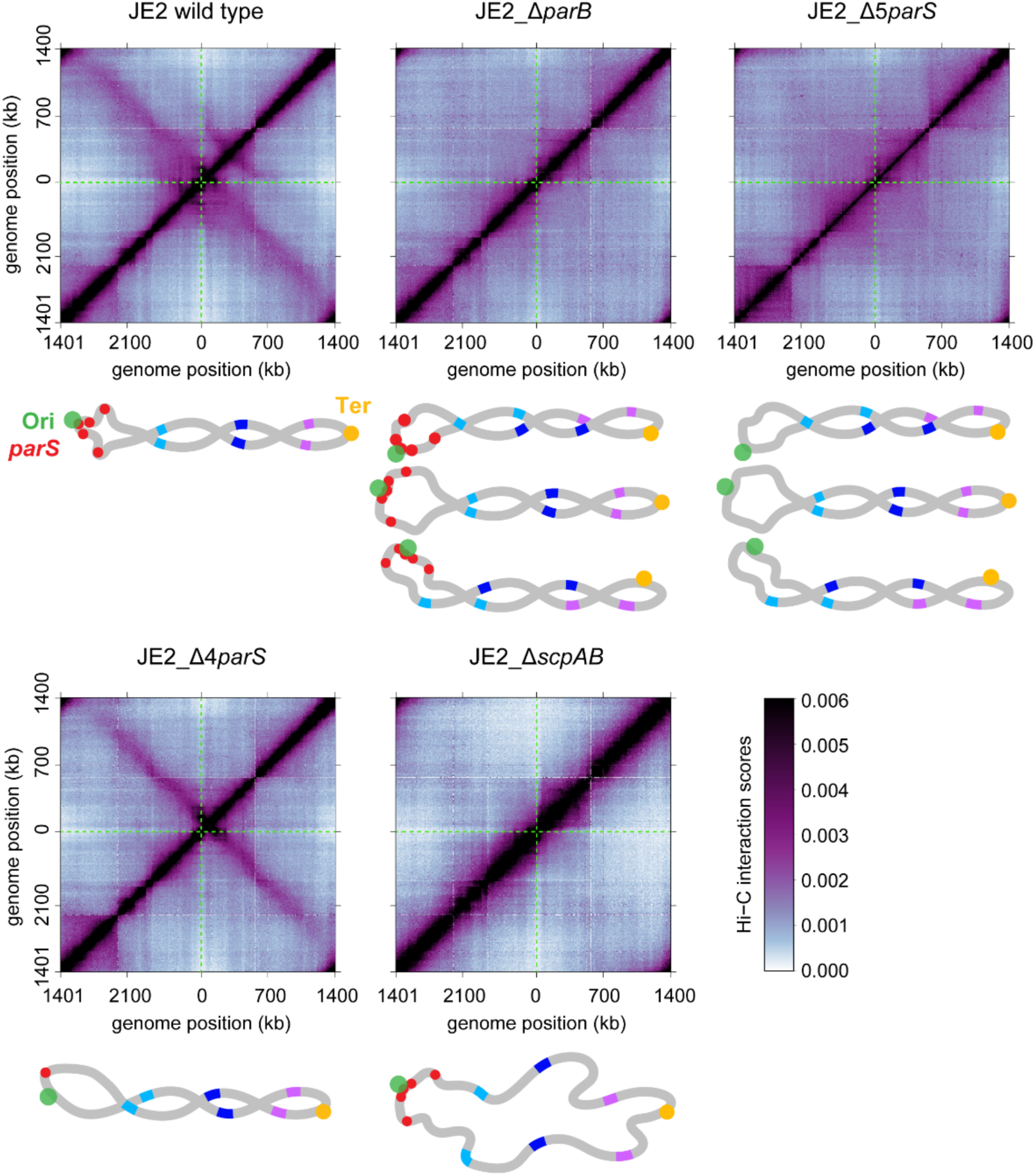
Role of ParB-*parS* and SMC-ScpAB in chromosome organization. Normalized Hi-C maps showing contact frequencies for pairs of genomic regions (5-kb bins) which are represented by the Hi-C score (lower right color scale) in the indicated strains. The chromosomal origin is at the center of the axes. The primary diagonal, running from lower-left to upper-right, represents local, short-range interactions, while the secondary diagonal, running upper-left to lower-right, represents interactions between loci equidistant from the Ori and on opposite chromosomal arms. Green dotted lines cross the maps horizontally and vertically, through the center (Ori), where the two diagonals intersect in JE2, but not in JE2_Δ4*parS*. Below each map a scheme shows the proposed chromosome arrangement, with the Ori in green, the *parS* sites in red and the Ter in yellow. Pairs of loci that, in JE2, would be on opposite arms and in physical proximity (long-range contacts), are represented as colored line segments. Note that the specific pattern of long-range contacts that results in the secondary diagonal in JE2, is lost in the JE2_Δ*parB* and JE2_Δ5*parS* strains. In these strains SMC is no longer loaded at specific *parS* sites, leading to different possible chromosome arrangements, three examples of which are represented.

To assess the role of the SMC complex in chromosome organization, we made a clean deletion of the genes *scpA* and *scpB* (JE2_Δ*scpAB*), whose inactivation in *B. subtilis* leads to the same phenotype as that of SMC (*74*). Similarly to deletion of *parB,* deletion of *scpAB* also abolished the secondary diagonal in the HiC map, consistent with the idea that ParB loading SMC complexes to *parS* sites generates arm alignments. In addition, deletion of *scpAB* caused a reduction in the Hi-C signal outside of the primary diagonal (i.e. long-range DNA interactions) and an increase in the signal of the primary diagonal (i.e. short-range DNA interactions), in comparison to both JE2_Δ*parB* and JE2 wild type (Fig. 3). Indeed, when we analyzed the global contact probability curve, Pc(s) curve, which shows the averaged contact probability for all loci separated by a set distance, we found that the JE2 and JE2_Δ*parB* had almost identical curves, but JE2_Δ*scpAB* showed reduced long-range DNA interaction (Fig. S5C). These results indicate that the SMC complex is responsible for forming DNA interactions between regions that are more than 400 kb apart, but this activity does not require specific loading of the SMC complex at *parS* sites.

To assess the role of ParB and the SMC complex in chromosome segregation, we examined the localization and number of Ori and Ter foci, as well as the occurrence of anucleate cells, in mutants lacking these proteins. Deletion of *parB* in the background of strains JE2_FROS^Ori^ and JE2_FROS^Ter^ did not alter the localization of Ori and Ter foci in comparison with the parental control strains (Fig. 4A), but led to a reduction in cells with four Ori foci (Fig. 4B). Importantly, the JE2_Δ*parB* strain produced only 0.14±0.04% of anucleate cells (vs. 0.03±0.05% in JE2 wild type, Fig. S6), consistent with previous reports suggesting that *parB* deletion causes only a very mild chromosome segregation defect in *S. aureus* (*63*, *75*).

**Figure 4.**
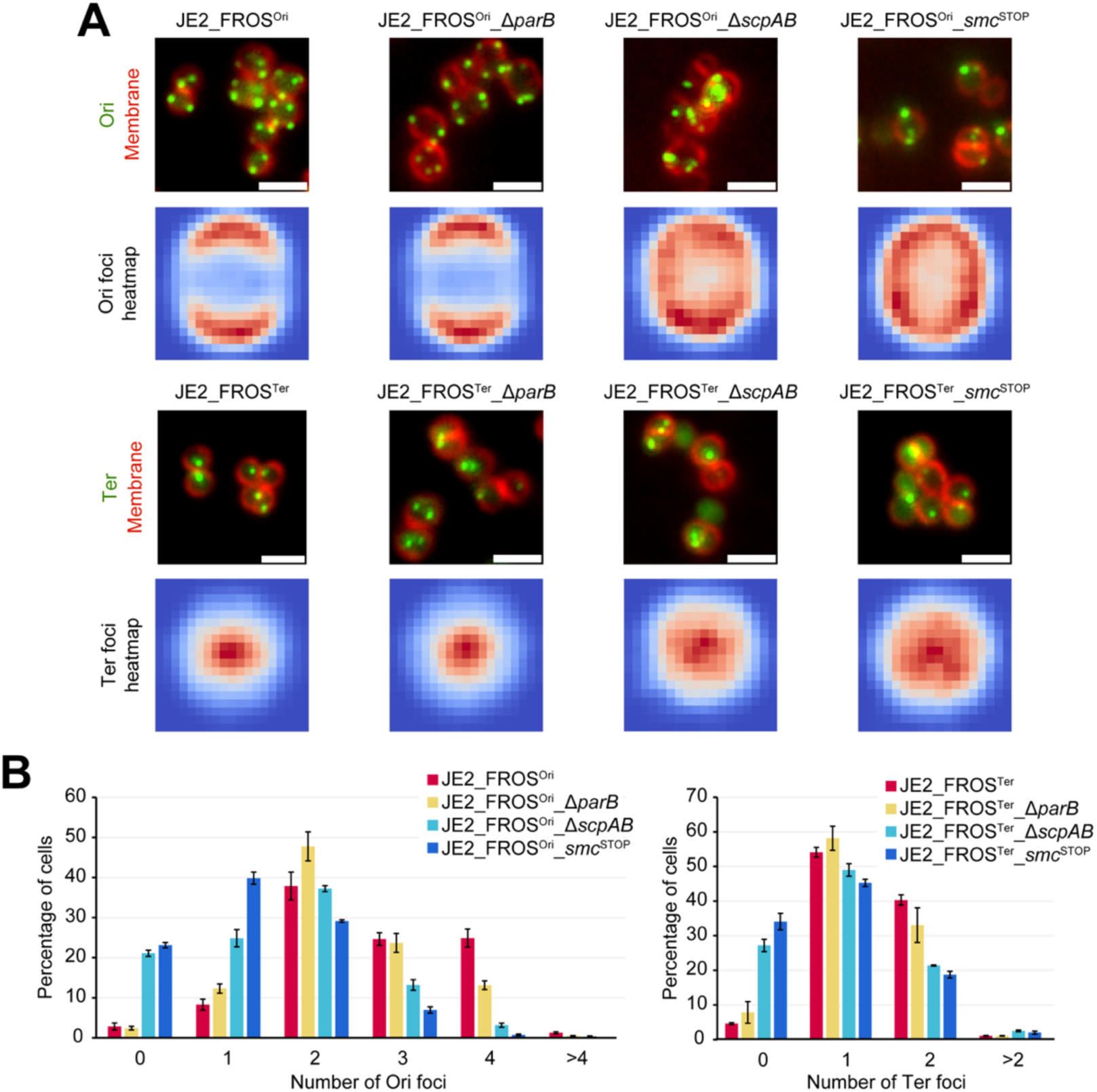
Role of ParB and SMC-ScpAB in chromosome segregation. Fluorescence microscopy images of the indicated strains showing localization of Ori (top) or Ter (bottom) labeled with the FROS system (green) and membrane labeled with FM 4-64 dye (red). Scale bar: 2μm. Heatmaps of the average localization of detected fluorescence spots of each strain are shown below each microscopy image. The color scale in each dataset ranges from red (maximum spot density) to dark blue (no spots detected), n>3000 per dataset. **B** Bar charts showing the relative distribution of the number of Ori (left, n>800) and Ter (right, n>900) foci in cells of strains shown in A. Data from three biological replicates, error bars indicate standard deviation.

The function of the SMC complex in chromosome segregation was evaluated using two different mutants: the JE2_Δ*scpAB* strain described above and a second mutant in which we introduced an array of premature STOP codons near the start of the *smc* gene (JE2_*smc*^STOP^ strain), preventing potential polar effects on the two essential genes (*ftsY* and *ffh*) downstream of *smc* (*76*). Both mutants exhibited ∼17% of anucleate cells (Fig. S6). In agreement with this increased number of anucleate cells, mutants lacking a functional SMC complex showed a higher number of cells with no Ori foci and a decrease in cells with 3 or 4 Ori foci when compared with the parental strain (Fig. 4B). The localization of origins in both Δ*scpAB* or *smc*^STOP^ backgrounds was altered, with a distribution less restricted to the cell poles, though still in close proximity to the membrane (Fig. 4A). The termini also showed a more diffuse pattern, although still positioned around the cell center. Altogether, our data indicate that the SMC complex plays a key role in the spatial organization and segregation of the *S. aureus* chromosome. We note that previous reports on the effect of deletion of *smc* had conflicting results, with anucleate cells frequencies varying from 1-2% (*63*, *77*) to 10% (*78*), while deletion of *scpB* led to the production of ∼14% of anucleate cells (*79*). The discrepancy might be due to the occurrence of suppressor mutations in previous mutants.

Finally, we constructed a double mutant lacking both *parB* and *scpAB* (JE2_*ΔparB_ΔscpAB*) and found that it was very similar to JE2_Δ*scpAB* in terms of anucleate cells frequency (Fig. S6). Together with the Hi-C data from the JE2_Δ*parB* and JE2_Δ*scpAB* strains (Fig. 3), this result suggests that in *S. aureus,* which is missing ParA, the main role of ParB is to load SMC complexes at the *parS* sites. Since Δ*parB* alone had little effect on chromosome segregation but *smc*^STOP^ and Δ*scpAB* each had a strong defect in chromosome segregation, we conclude that the SMC complex does not need to be specifically loaded onto the *parS* sites to segregate chromosomes; without ParB/*parS*, randomly loaded SMC-ScpAB molecules can compact and segregate the chromosome quite well, although the overall organization of the chromosome is altered (Fig. 3).

### Role of factors connecting the divisome with the chromosome

One important link between the cell division machinery and chromosome segregation is the FtsK protein family of DNA pumps that ensure segregation of the chromosomes before the completion of the division septum (*80*). In *E. coli,* FtsK works together with the recombinase XerCD complex to resolve chromosome dimers that would otherwise prevent proper chromosome segregation (*81*). *S. aureus* has two FtsK family proteins, FtsK and SpoIIIE. Each protein is individually dispensable, but the presence of at least one is required for correct chromosome segregation (*82*). Visualization of the Ori and Ter regions in mutants lacking FtsK or SpoIIIE revealed a moderate reduction in the number of cells with four Ori foci (Fig. 5A, B), decreasing from 25% in the JE2_FROS^Ori^ strain to 19% in the Δ*ftsK* mutant and 15% in the Δ*spoIIIE* mutant. However, the two mutants had opposite phenotypes regarding the number of terminus foci (Fig. 5A, C). The JE2_FROS^Ter^_Δ*ftsK* mutant produced about 10% of cells with more than two termini, compared to only 1% in the JE2_FROS^Ter^ parental strain. This could be a consequence of the delay in cell division caused by the deletion of *ftsK* which, in *S. aureus*, leads to a longer cell cycle Phase 3 (*83*) allowing chromosome replication time to complete before cell division. On the other hand, the JE2_FROS^Ter^_Δ*spoIIIE* strain produced around 17% of cells with no terminus foci (compared to 4.5% in the JE2_FROS^Ter^ parental strain). A similar, but more pronounced, phenotype was observed in the JE2_FROS^Ter^_Δ*xerC* strain, with 35% of cells lacking a terminus focus. Marker frequency analysis, used to examine the relative abundance of specific DNA sequences across the genome, revealed a decreased DNA copy number at the terminus region in the population of Δ*xerC* mutant cells (Fig. S7, blue arrows). This suggests that the absence of XerC results in degradation of the terminus region (where the *tetO* array is located), preventing the formation TetR-mNG foci in the affected cells. Interestingly, marker frequency analysis did not show an obvious decrease in the terminus region in the JE2_FROS^Ter^_Δ*spoIIIE* strain, indicating that another process may be interfering with the TetR association to the *tetO* arrays in this strain (and potentially also in the JE2_FROS^Ter^_Δ*xerC* strain). Furthermore, in Δ*xerC* and Δ*spoIIIE* mutants, we observed two peaks, one in each replication arm, indicating increased DNA amplification in these genomic regions (Fig. S7, red arrows). These regions encompass genes that are annotated as encoding phage proteins, including capsid proteins, phage terminases and phage DNA primases (genes *SAUSA300_1921-1940* in the left arm peak and *SAUSA300_0809-0815* in the right arm peak). Therefore, it is likely that the observed DNA amplification in these regions is due to the activation of prophages, triggered by the deletion of *xerC* or *spoIIIE*.

**Figure 5.**
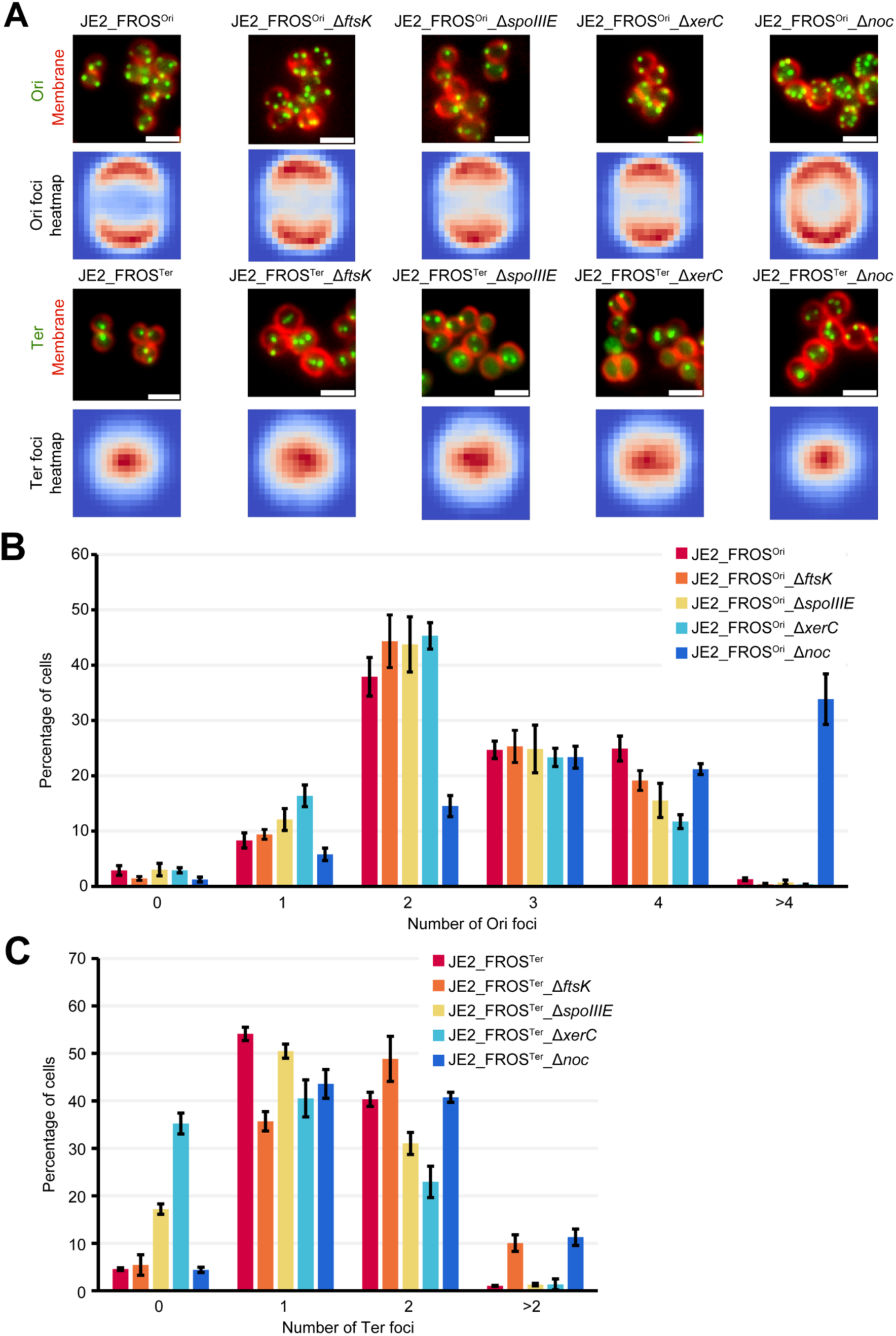
Impact of FtsK, SpoIIIE, XerC and Noc absence on Ori and Ter copy numbers. **A** Fluorescence microscopy images of the indicated strains showing localization of Ori or Ter labeled with the FROS system (green) and membrane labeled with FM 4-64 dye (red). Scale bar: 2μm. Heatmaps of the average localization of detected fluorescence spots of each strain are shown below each microscopy image. The color scale in each dataset ranges from red (maximum spot density) to dark blue (no spots detected), n>2500 each. **B** Bar chart showing the relative distribution of number of origin foci in cells of the strains shown in A. Data from three biological replicates, error bars indicate standard deviation, n>500. **C** Bar chart showing the relative distribution of number of terminus foci in cells of the strains shown in A. Data from three biological replicates, error bars indicate standard deviation, n>800.

Despite the changes in the number of origins and termini in mutants lacking *ftsK, spoIIIE* or *xerC,* the cellular localization of Ori and Ter foci remained similar suggesting that these proteins act on chromosome segregation but do not have a major role in the spatial placement of chromosome loci.

We also investigated the role of the nucleoid occlusion protein Noc, which in both *B. subtilis* and *S. aureus* prevents the assembly of the divisome over the nucleoid to avoid its guillotining (*47*, *49*). Additionally, in *S. aureus,* Noc is a negative regulator of the initiation of DNA replication (*75*). In agreement with published data, the JE2_FROS^Ori^_Δ*noc* strain exhibited over 30% of cells with more than four origin foci (Fig. 5A, B), and showed an increased number of cells with more than two terminus foci (Fig. 5C). Despite the increased number, the Ori foci remain spaced from each other, although they were more dispersed around the cell periphery compared to the JE2_FROS^Ori^ parental strain (Fig. 5A). Overall, our data supports the role of Noc as a key regulator of chromosome replication in *S. aureus*.

## Discussion

The spatial and temporal organization of the bacterial chromosome may seem particularly challenging for a nearly spherical bacterium, as there are fewer geometric cues available compared to rod-shaped or asymmetric cells. Yet, *S. aureus* elegantly solves this problem for chromosome segregation, by becoming a pseudo-diploid and decoupling its chromosome replication cycle (one complete round of chromosome replication) from its cell division cycle (from completion of one cell division to the next). *S. aureus* newborn cells, in fast growing conditions, typically have two origins of replication and one active replisome (Fig. 6A), similar to *B. subtilis* in slow growing media (*6*). The origins tend to localize at the cell periphery, near the membrane, positioned opposite to each other, at the celĺs poles. This arrangement establishes an axis of chromosome segregation, breaking the internal spherical symmetry of the cell. As chromosomes segregate along this axis, a DNA free region is generated between them, providing space for a division site, where the septum can form without guillotining the DNA (Fig. 6B). Mechanistically, this is mediated by the nucleoid occlusion protein Noc, which binds the origin-proximal region of the chromosome and prevents spurious FtsZ assembly in those regions (*47*). A second round of replication can begin in Phase 1 cells, i.e., even before septum synthesis starts. This is supported by the observation that ∼35% of Phase 1 cells have three to four segregated origins (Fig. 1B), indicating that the presence of a septum is not required for origin segregation. As the cell cycle progresses and the septum begins to be synthesized (Fig. 6B), the cell becomes increasingly divided in two hemispherical compartments. These compartments now have a long axis (parallel to the nascent septum) and a short axis (perpendicular to the nascent septum). Our data show that, within each hemisphere, the two origins generally segregate away from each other along a long axis parallel to the septum (Fig. 6C). Therefore, when P3 cells split and give rise to newborn P1 cells, the future division plane is already defined within the spherical cytoplasmic compartment, located between the segregated chromosomes, where the septum will form. In turn, the septum demarcates the possible directions for the next round of chromosome segregation.

**Figure 6.**
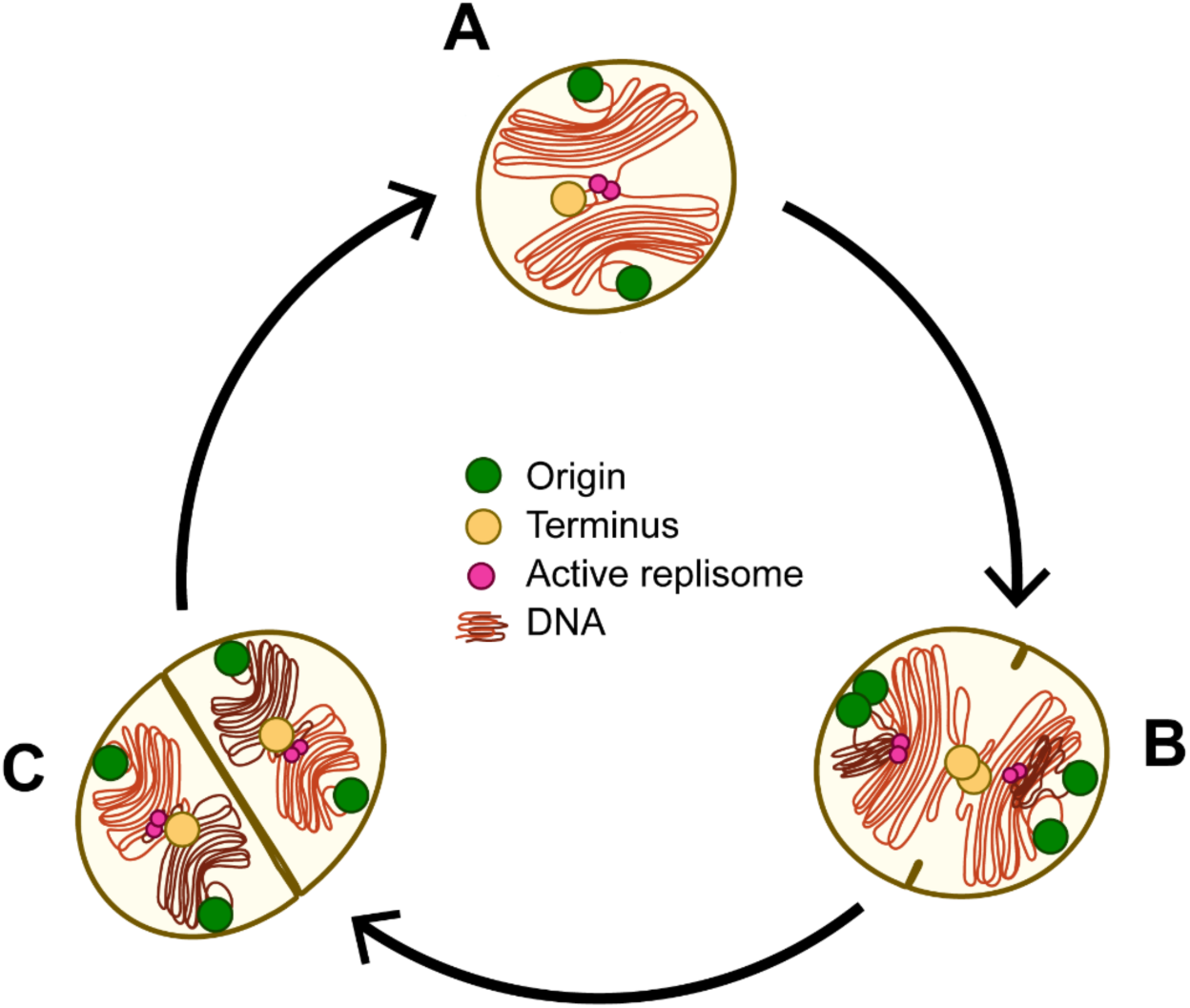
Representation of chromosome organization and dynamics of a typical *S. aureus* cell cycle. The cell envelope (brown), origins (green circles), active replisomes (magenta circles), termini (yellow circles) and the chromosome (orange and brown) are illustrated. **A** Typical newborn cell (P1), with two segregated origins and a hemi-replicated chromosome. **B** P2 cell, after the new round of replication has begun, and origins segregation has initiated. **C** P3 cell, which generally has four segregated origins and each hemi-replicated chromosome occupying one of the hemispheres, which become spherical again after cell division. Notice that the cell cycle and chromosome replication cycle are not coupled and Ori segregation can occur as early as P1, leading to ∼35% of P1 cells having three or four origins.

Some fast-growing organisms, such as *B. subtilis* (*15*) and *E. coli* (*16*), as well as slow-growing like *M. smegmatis* (*17*) can undergo multifork replication which occurs when multiple rounds of replication take place during one cell cycle, usually under rich media conditions. However, we did not detect *S. aureus* cells with assembled replisomes colocalizing with the origins while a second set of replisomes was located further away from the origins. Combined with the absence of newborn cells exhibiting more than two origin foci, this indicates that multifork replication is not typical in *S. aureus* cells under the tested fast-growth conditions. Coincidentally, the ovococcal *S. pneumoniae* is also thought not to engage in multifork replication (*13*). It is possible that the small size and the geometry of *S. aureus* (and other coccoid bacteria) makes it difficult to coordinate multiple rounds of replication in a single cell cycle.

The chromosomal origin in *S. aureus* is primarily localized at the cell periphery, near the membrane, throughout the cell cycle (Fig. 2C, E), even in mutants affecting its number (Δ*noc,* Fig. 5A) or segregation (Δ*scpAB*, *smc*^STOP^, Fig. 4A). This localization is not exclusive of *S. aureus*, as other bacteria also position their origins in close proximity to the membrane, such as *B. subtilis* during sporulation or *S. pneumoniae* (*84*, *85*), or in polar or sub-polar regions, like *C. crescentus*, or *M. xanthus* (*3*, *4*). These bacteria employ molecular mechanisms to restrict the movement of the origins, leading us to hypothesize that *S. aureus* likely has a similar mechanism, perhaps similar to those in other firmicutes, such as the RacA protein in *B. subtilis* (*85–87*) or RocS in *S. pneumoniae* (*84*).

The forces driving chromosomal segregation in *S. aureus* are not yet fully understood, but our data from the Δ*scpAB* and *smc*^STOP^ mutants strongly suggest that the SMC complex plays a crucial role in the process, perhaps by promoting the unmixing of the sister chromosomes as they are replicated (*29*). Furthermore, the steep increase in cells with a single Ori focus in the Δ*scpAB* and *smc*^STOP^ mutants (Fig. 4B) suggests that the SMC complex has an important role in the segregation of the Ori regions. This is in line with the drastic decrease in cells with three or four origins in these mutants, compatible with the possibility that chromosomes are replicated but origins remain together and are thus indistinguishable using the FROS system. Such a failure in origin segregation would compromise the overall chromosome segregation process, resulting in the production of anucleate cells, as observed (Fig. S6). Importantly, ParB is dispensable for overall chromosome segregation in *S. aureus*, although its presence organizes the loading of the SMC complexes (Fig. 3). In the Δ*parB* background, we measured a reduction in cells with four Ori foci, suggesting that ParB-mediated loading of the SMC complexes slightly increases the efficiency or speed of Ori segregation. Overall, our findings indicate that ParB binding to *parS* loads SMC, which globally organizes the chromosome, creating a defined inter-arm alignment. However, this specific alignment by itself is not a major contributor to chromosome segregation. Rather, SMC loading onto the chromosome (not necessarily at *parS* sites) and its translocation away from the loading position, generates DNA loops (long-range DNA contacts), which simultaneously compacts the chromosomes and promotes their segregation.

Two additional proteins involved in chromosome segregation in *S. aureus* are the DNA pumps FtsK and SpoIIIE. Each protein is individually dispensable, but at least one must be present for correct chromosome segregation, suggesting partial redundancy (*82*). However, FtsK and SpoIIIE do not colocalize, and while *ftsK* deletion causes cell morphology defects such as multi-septated cells and cell size heterogenicity, deleting *spoIIIE* leads to an increase of cells with condensed chromosomes, altogether indicating that they have partially independent functions (*82*). In this study, deletion of *ftsK* led to an increase in the number of Ter foci (Fig. 5C). Previous research has shown that FtsK mutants have a delay in P3, resulting in cells remaining for longer in a pre-divisional stage. This delay is due to an additional role of *S. aureus* FtsK in promoting the export of the autolysin Sle1, a peptidoglycan hydrolase that plays an important role in splitting the septum at the end of the cell cycle (*83*). Therefore, it is plausible during this delay, chromosome replication has time to complete, explaining the observed increase in the number of Ter foci. In contrast, deletion of *spoIIIE* results in approximately 17% of cells lacking a terminus focus, a phenotype similar to that observed in the Δ*xerC* mutant. In *E. coli,* the XerC recombinase works together with FtsK to resolve chromosome dimers during the final stages of chromosome replication and segregation (*81*, *88*). In *S. aureus,* previous studies have shown that deleting either *spoIIIE* or *xerC* increases the number of cells with condensed nucleoids, with this effect being more pronounced in the Δ*xerC* mutant (*82*). However, we found no correlation between cells with condensed nucleoids and cells lacking Ter foci. Furthermore, we also showed that deletion of *xerC* results in degradation of the Ter region, potentially explaining why a subset of cells lack a Ter focus (Fig. S7). However, the literature presents conflicting evidence regarding Ter degradation in *E. coli xerC* mutants. While one study reports Ter degradation (*89*), another finds minimal differences compared to the wild-type strain (*90*). Collectively our findings serve as a starting point for further investigation into possible functional connections between SpoIIIE and XerC, a link suggested by a previous study (*82*).

Finally, our data supports the proposed role of Noc as a key regulator of initiation of DNA replication (*75*), given that its absence led to a sharp increase in the number of cells with more than four origins.

This study provides the first comprehensive characterization of chromosome positioning and dynamics in a small, spherical bacterium, highlighting the role of chromosome segregation in division site positioning. When comparing to other studied organisms, *S. aureus* chromosome organization and replication cycle resembles that of slow-growing *B. subtilis*, where newborn cells typically start with one hemi-replicated chromosome and origins positioned at opposite poles. However, key differences were observed in *S. aureus*, such as origin being consistently associated with the cell periphery and segregation occurring along an axis parallel to the septum. Future research will determine whether other spherical coccoid organisms follow a similar pattern.

## Materials and Methods

### Bacterial growth conditions

Strains of *E. coli* were grown in lysogeny broth (LB, VWR) or on lysogeny broth agar (LA, VWR) at 37°C. *S. aureus* was grown in tryptic soy broth (TSB, Difco) with agitation (200 rpm) or on tryptic soy broth agar (TSA, VWR). When required, media were supplemented with antibiotics (100 μg mL^−1^ ampicillin, Sigma-Aldrich; 10 μg mL^−1^ erythromycin, Apollo Scientific). For blue/white colony screening, TSA plates were supplemented with 5-bromo-4-chloro-3-indolyl β-d-galactopyranoside (X-Gal, Apollo Scientific) at 100 μg mL^−1^. When required, Cadmium chloride (Fluka) was added to liquid cultures at 1 µM and Isopropyl β-D-1-thiogalactopyranoside (IPTG, NZYtech) added at 100 µM.

### Plasmid and strain construction

The complete lists of strains, plasmids and oligonucleotides are in Supplementary Table 1, 2, and 3, respectively. Plasmids were assembled as described in Supplementary Table 2, propagated in *E. coli* DC10B and purified using the QIAprep Spin miniprep kit (Qiagen) and verified by sequencing. Purified plasmids were used to transform by electroporation *S. aureus* RN4220 cells as previously described (*91*) and subsequently transduced into *S. aureus* JE2 or strains in this background using the bacteriophage 80α(*92*). *S. aureus* strain construction was done using derivatives of the temperature-sensitive vector pMAD (*93*), indicated in Supplementary Table 1, by performing allelic replacement through double homologous recombination, creating marker-less strains. Allelic replacement was confirmed by colony polymerase chain reaction (PCR).

### Molecular biology methods

Amplification of DNA fragments for plasmid construction was carried out using a Phusion high-fidelity polymerase kit (Thermo Scientific) following the manufacturer instructions. For PCR using as a template a *S. aureus* bacterial colony, a small portion of the colony was resuspended in phosphate buffered saline (PBS; 137 mM NaCl, 2.7 mM KCl, 10 mM Na_2_HPO_4_, 1.8 mM KH_2_PO_4_) and cells were disrupted mechanically (by adding glass sand and three cycles of shaking for 45s at a speed of 6.5 m s^−1^ in a FastPrep-24, MP Biomedicals) or enzymatically (by incubation at 37°C for 1 h in the presence of 10 µg mL^−1^ of lysostaphin, Sigma) and the lysate was used as PCR template. For the PCR reaction, the Phire Hot Start II PCR Master Mix (Thermo Scientific) was employed following the manufacturer instructions.

Cloning was performed using restriction enzymes (FastDigest, Thermo Scientific) indicated, for each construct, in Supplementary Table 2. Fragments were ligated using T4 DNA ligase (Thermo Scientific). For Gibson Assembly, the Gibson Assembly Master Mix (NEB) was employed.

### Microscopy

*S. aureus* strains were streaked from cryo-stocks onto TSA plates. Single colonies were used to inoculate independent cultures in TSB that were grown overnight at 37°C with agitation. The next day, the cultures were diluted 1:200 in TSB and grown at 37°C with agitation until they reached mid-exponential phase (OD_600_ 0.6-0.8). Fluorescent dyes for membrane labelling (FM4-64, 5 µg mL^−1^, Invitrogen; CellBrite Fix 640 3.3 nM, Biotium), DNA labelling (Hoechst 33342, 1 µg mL^−1^, Invitrogen) or HaloTag (HT) labelling (Janelia Fluor 549 HT ligand, 500 nM, Janelia Research Campus) were added when required to 1 mL of the exponential culture, which was then incubated for 20 minutes at 37°C with agitation. Afterwards, the culture was centrifuged at 10000 ×g for one minute and the pellet was resuspended in 50 µL of PBS and 1 µL of the suspension was spotted on a pad of 1.2% Topvision Agarose (Thermo Fisher) in PBS.

For time-lapse microscopy, the aforementioned procedure was followed, but cells were spotted on pads of 1.2% Topvision Agarose (Thermo Fisher) in M9 minimal medium (KH_2_PO_4_ 3.4 g L^−1^, VWR; K_2_HPO_4_ 2.9 g L ^−1^, VWR; di-ammonium citrate 0.7 g L^−1^, Sigma-Aldrich; sodium acetate 0.26 g L^−1^, Merck; glucose 1% (w/v), Merck; MgSO_4_ 0.7 mg L^−1^, Sigma-Aldrich; CaCl_2_ 7 mg L^−1^, Sigma-Aldrich; casamino acids 1% (w/v), Difco; minimum essential medium amino acids 1×, Thermo Fisher Scientific; and minimum essential medium vitamins 1×, Thermo Fisher Scientific). The cells were kept at 37°C during the imaging procedure and were imaged every three minutes.

Imaging was performed in a DeltaVision OMX SR microscope equipped with a hardware-based focus stability (HW UltimateFocus) and an environmental control module (set to 37° for time-lapses). *Z*-stacks of three epifluorescence images with a step size of 500 nm were acquired using a 405 nm laser (100 mW, at 10% maximal power; for the Hoechst 33342 DNA dye), a 488 nm laser (100 mW, at 15% maximal power for the mNeonGreen fusions), a 568 nm laser (100 mW, at 30% maximal power; for JF549-labelled DnaN-Halo and FM4-64 membrane dye) or a 640 nm laser (100 mW, at 40% maximal power; for the CellBrite Fix 640 dye), each with an exposure time of 100 ms. When required, a maximum intensity projection of the three images from each *Z*-stack, fluorescence channel alignment and SIM image reconstruction was performed using SoftWoRx v7.2.1.

For cell cycle automated classification (Fig. 2E), cells were imaged in a Zeiss Axio Observer microscope equipped with a Plan-Apochromat 100×/1.4 oil Ph3 objective, a Retiga R1 CCD camera (QImaging), a white-light source HXP 120 V (Zeiss) and the software ZEN blue v2.0.0.0 (Zeiss). For image acquisition, the filters (Semrock USA) Brightline TXRED-4040B (FM4-64), Brightline GFP-3035B (mNeonGreen), and Brightline DAPI-1160A (Hoechst 33342) were used.

### Image processing and automated analysis

Images were examined using ImageJ Fiji (*94*), which was also used to produce crops of illustrative regions. Lateral drift in time-lapse datasets was corrected with the ImageJ plugin NanoJ (*95*).

For cell cycle automated classification and generation of foci average heatmaps from images obtained using the Zeiss Axio Observer microscope (figure 2E), crops of single cells and automated cell cycle phase analysis were generated using eHooke software version 1.1, as previously described (*70*).

For foci quantification and generation of foci average heatmap from images acquired in the OMX microscope (figures 2C, 2D, 4A, 4B, 5A, 5B, 5C), cell segmentation was performed using an in-house fine-tuned StarDist model (*96*) applied on images with fluorescence signal from membrane labelling. When mentioned, the cell cycle phase analysis was performed manually (figure 2D).

After cell segmentation (and cell cycle classification if required) we used a PCA transform applied to the coordinates of the pixels that constitute the cell outline to calculate the orientation of the major axis of each cell. Then cell crops were aligned by their major axes as previously described (*83*). Foci localization was determined using TrackMate 7.11.1 (*71*) using the Laplacian of Gaussian filter with subpixel localization. The blob diameter was set to 0.24 μm, the quality threshold was manually adjusted for each field of view, and the results were exported as a .xml file. In each cell crop, foci were represented as a circle of 1 pixel radius, intensity 1, and the same relative coordinates as the foci, in a rectangle with the same dimensions as the cell crop (model image), with background set to 0. All model images were then resized to a common width and height equal to the median of the width and height of all cell crops. Heatmaps were generated by averaging all model images, and colored using the coolwarm colormap provided by matplotlib (*97*).

### High-throughput Chromosome Conformation Capture (Hi-C)

The Hi-C procedure used for *S. aureus* was adapted from a previously described protocol used for *B. subtilis* (*98*, *99*). Briefly, *S. aureus* strains were streaked from cryo-stocks onto TSA plates. Single colonies were used to inoculate independent cultures in TSB that were grown overnight at 37°C with agitation. The next day, the cultures were diluted 1:1000 in TSB and grown at 37°C with agitation until they reached early-exponential phase (OD_600_ 0.3-0.4). Cells were crosslinked by adding formaldehyde (Sigma) to a final concentration of 7% (v/v) at room temperature (RT) for 30 min and quenched with 125 mM glycine (Sigma). Cells were lysed using Ready-Lyse Lysozyme (Epicentre, R1802M) and 200 µg mL^−1^ lysostaphin (Sigma-Aldrich, L9043) at RT for 1 h, followed by the treatment with 1% SDS (v/v) at RT for 30 min. Solubilized chromatin was digested with DpnII for two hours at 37°C. The digested ends were filled in with Klenow and Biotin-14-dATP, dGTP, dCTP, dTTP. The products were ligated with T4 DNA ligase at 16°C for about 20 h. Crosslinks were reversed at 65° C for 17-20 h in the presence of EDTA, proteinase K and 0.5% SDS. The DNA was then extracted twice with phenol/chloroform/isoamylalcohol (25:24:1) (PCI), precipitated with ethanol, and resuspended in 20 µL of 0.1X TE buffer (10 mM Tris-HCl, 1 mM EDTA). Biotin from non-ligated ends was removed using T4 polymerase (4 h at 20°C) followed by extraction with PCI. The DNA was then sheared by sonication for 12 min with 20% amplitude using a Qsonica Q800R2 water bath sonicator. The sheared DNA was used for library preparation with the NEBNext UltraII kit (E7645). Biotinylated DNA fragments were purified using 5 µL streptavidin beads. DNA-bound beads were used for PCR in a 50 µL reaction for 14 cycles. PCR products were purified using Ampure beads (Beckman, A63881) and sequenced at the Indiana University Center for Genomics and Bioinformatics using NextSeq500. Paired-end sequencing reads were mapped to the genome of *S. aureus* JE2 (NCBI Reference Sequence GCF_002085525.1) using the same pipeline described previously (*99*). The genome was divided into 5-kb bins. Subsequent analysis and visualization were done using R scripts. Hi-C scores, which quantify the interaction between loci and correct for biases in the abundance of the different bins in each experiment, were calculated as described in (*99*).

### Whole Genome Sequencing (WGS)

For genomic DNA extraction, cells from the relevant strains were grown in TSB at 37°C with agitation overnight. The next day the cultures were diluted 1:200 in 50 mL of TSB and grown at 37°C with agitation until the mid-exponential phase (OD_600_ 0.6-0.8). Then cultures were centrifuged at 6000 ×g for 10 min and the supernatants were discarded. Cells were resuspended in 180 µL of Enzymatic lysis buffer (20 mM TRIS, VWR; 2 mM sodium EDTA; 1.2% (v/v) Triton X-100, Sigma, adjusted to pH 8 using HCl, Sigma), supplemented with 100 µg mL^−1^ of lysostaphin (Sigma) and were incubated at 37°C for 15 min. Afterwards, the samples were processed using the DNeasy Blood & Tissue Kit (Quiagen) following the indications from the manufacturer. The extracted DNA was sonicated using a Qsonica Q800R2 water bath sonicator, prepared using the NEBNext UltraII kit (E7645), and sequenced at the Indiana University Center for Genomics and Bioinformatics using NextSeq500. The reads were mapped to the genome of *S. aureus* JE2 (NCBI Reference Sequence GCF_002085525.1) using CLC Genomics Workbench (CLC Bio, QIAGEN). The mapped reads were normalized by the total number of reads. Plotting and analysis were performed using R scripts.

### Chromatin immunoprecipitation (ChIP-seq)

The *S. aureus* strains JE2_ParB-3xFLAG and JE2_3xFLAG-mNG were grown overnight in TSB at 37°C with agitation. The next day the cultures were diluted 1:200 in 50 mL of TSB and incubated at 37°C with agitation until they reached the mid-exponential phase (OD_600_ 0.6-0.8). Cultures of the strain JE2_3xFLAG-mNG were supplemented with 100 µM IPTG to induce expression of *3xflag-mng.* Formaldehyde (Sigma) was added to a final concentration of 1% (v/v) and the mixture was incubated at room-temperature with shaking for 30 min. Afterwards, glycine was added to a final concentration of 125 mM and cultures were further incubated at room-temperature with shaking for 10 min. The mixture was cooled down on ice and centrifuged at 7000 ×g for 10 min at 4°C. The pellet was resuspended in ice-cold PBS and centrifuged again as in the previous step. This was repeated three times, before snap-freezing the pellet in liquid nitrogen and storing it at −80°C. When required, samples were thawed, resuspended in 300µl IP buffer (50 mM Tris/HCl pH 7.5, 5 mM EDTA, 150 mM NaCl, 25 mM sucrose, 1 µg mL^-1^ lysostaphin, Sigma, 0.3 µg mL^-1^ RNase A, Sigma, and 1 tablet of cOmplete protease inhibitor cocktail, EDTA free, Roche, per 10 mL of buffer) and incubated at 37°C for 1h with shaking. Afterwards the samples were cooled on ice, followed by addition of TritonX-100 to a final concentration of 1% (v/v). Samples were then sonicated using a Bioruptor Plus bath sonicator at 4°C using 50 cycles of alternating 30s on and 90s off in the high-power mode, followed by centrifugation at 20000 ×g for 10 min at 4°C. The supernatants were mixed with 50 µL of anti-Flag M2 agarose beads (Sigma, pre-washed in 1mL of IP buffer supplemented with 1% Triton X-100), and the mixture was incubated overnight at 4°C with tumbling. Afterwards, the IP-samples were centrifuged at 800 ×g for 2 min at 4°C, the supernatant was discarded, and the beads were resuspended in 1 mL of IP buffer with 1% Triton X-100. The IP-samples were centrifuged as in the previous step and resuspended in 1 mL of High-Salt Buffer (50 mM Tris/HCl pH 7.5, 5 mM EDTA, 700 mM NaCl, 0.1% Na-deoxycholate, Calbiochem, 1% Triton X-100). IP-samples were centrifuged as in the previous step and resuspended in 1 mM of TE buffer (10 mM Tris/HCl pH 8, 1 mM EDTA), this step was done twice. Then, the IP-samples were centrifuged and resuspended in 300 µL of Reversal Buffer (RB, 10 mM Tris/HCl pH 8, 1 mM EDTA, 300 mM NaCl), followed by addition of SDS to a final concentration of 1% (w/v). All samples were then incubated at 65°C with 1500 rpm shaking for 14-16 h. Afterwards, the IP-samples were centrifuged at 800 ×g for 2 min and the supernatant was transferred to a new tube. Then 300 µL of phenol-chloroform-isoamyl alcohol mix (Roth) were added to each sample, the mixture was vigorously mixed by vortexing for 10 s and centrifuged at 20000 ×g for 5 min at RT. 250 µL were taken from the aqueous phase and transferred to a new tube where they were combined with 25 μL of 3 M sodium-acetate (pH 5.2), 1.5 μL of 20 mg mL^−1^ glycogen and 690 μl of absolute ethanol. The samples were placed at −80°C for 1 h and then centrifuged at 20000 ×g for 15 min at RT. The supernatant was discarded and the pellet was washed with 1mL of ice-cold 70% (v/v) ethanol. Samples were centrifuged at 20000 ×g for 1 min at RT and the supernatant was discarded. The pellet was left to air-dry. Afterwards the pellet was resuspended in 25μl of Nuclease-free water and incubated at 55°C for 10 min with gentle shaking. Samples were sent to Lausanne Genomic Technologies Facility for next-generation sequencing. Sequencing results were assembled to the reference genome (NCBI Reference Sequence GCF_002085525.1) using the CLC workbench (Qiagen) and plotted using Microsoft Excel.

## Acknowledgements

We thank members of the Pinho lab, P. Pereira (ITQB-NOVA) and S. Filipe (FCT-NOVA) for stimulating discussions and support, H. Veiga (ITQB-NOVA) for reviewing the manuscript, N. Reichmann (ITQB-NOVA) for constructing plasmid pBCB33, N. Meiresonne and T. den Blaauwen (Univ of Amsterdam) for plasmid psav-mSc-I, L. Lavis (Janelia Research Campus, Ashburn) for the generous gift of JF549-HTL, and the Indiana University Center for Genomics and Bioinformatics for high throughput sequencing. This study was funded by the European Research Council through ERC Advanced Grant 101096393 (to MGP), by La Caixa Foundation grant HR23-00221 (to MGP), by Fundação para a Ciência e a Tecnologia (FCT) through grant 2022.01678.PTDC (to SS) and contract 2022.03033.CEECIND (to SS), MOSTMICRO-ITQB R&D Unit (UIDB/04612/2020, UIDP/04612/2020 to ITQB-NOVA) and LS4FUTURE Associated Laboratory (LA/P/0087/2020 to ITQB-NOVA), and by National Institutes of Health R01GM141242, R01GM143182, and R01AI172822 (to XW). This research is a contribution of the GEMS Biology Integration Institute, funded by the National Science Foundation DBI Biology Integration Institutes Program, Award #2022049 (XW).

## Data and code availability

The codes used to create average localization heatmaps were deposited to github (https://github.com/BacterialCellBiologyLab/AverageCellLoc/releases/tag/1.0.0).

## Supplementary Figures and Tables

**Supplementary Figure 1.**
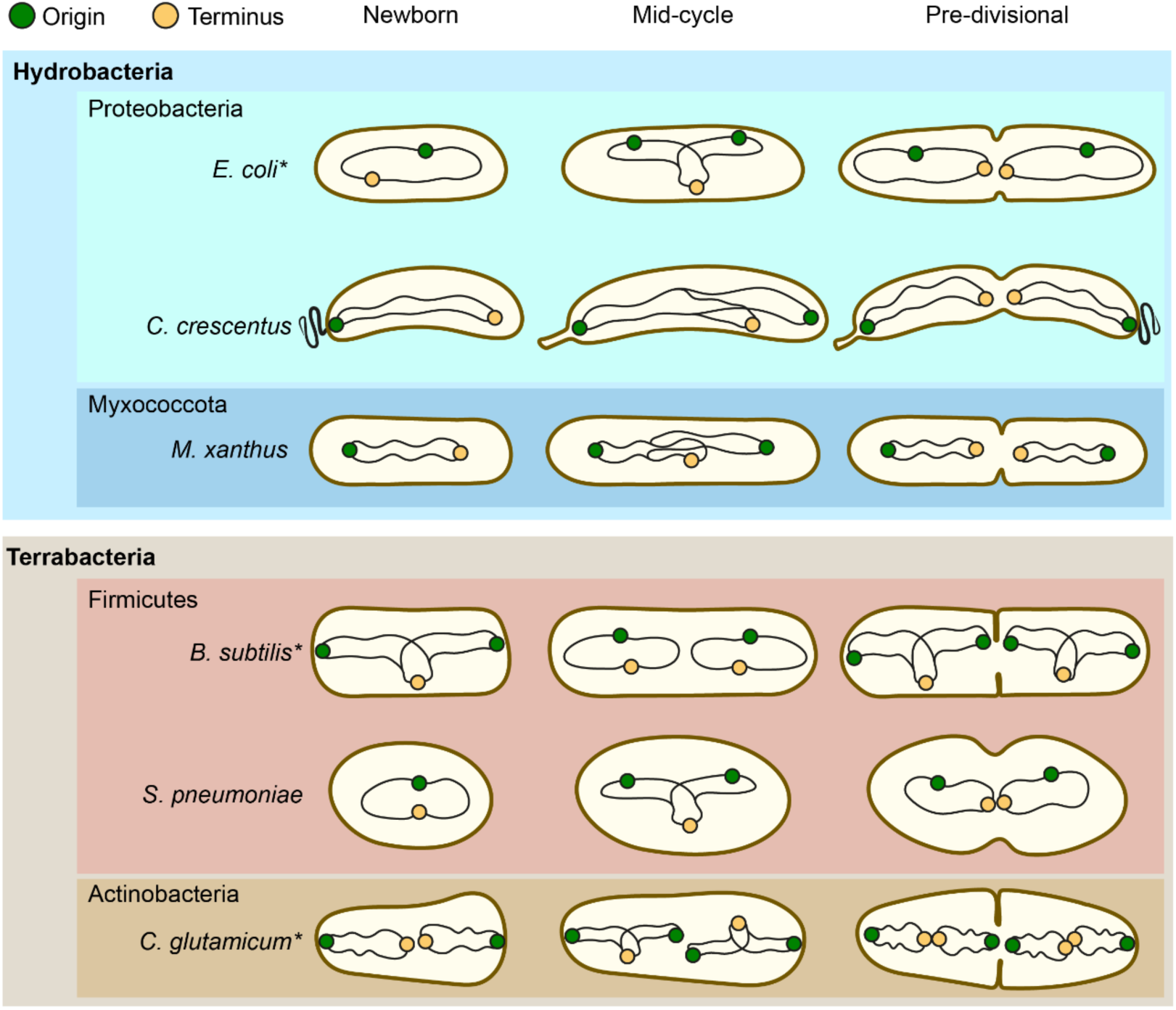
Chromosome organization and dynamics in various bacteria. Chromosome are depicted as black lines, with the origin and terminus as green and yellow dots, respectively. Species are arranged according to their taxonomy, as indicated by the colored boxes labelled with the corresponding phyla. Cells are represented, from left to right, as newborn, in an intermediate stage and pre-divisional. For species marked with an asterisk, the diagram represents chromosome dynamics under slow growing conditions (in the absence of multifork replication). References are provided in the main text.

**Supplementary Figure 2.**
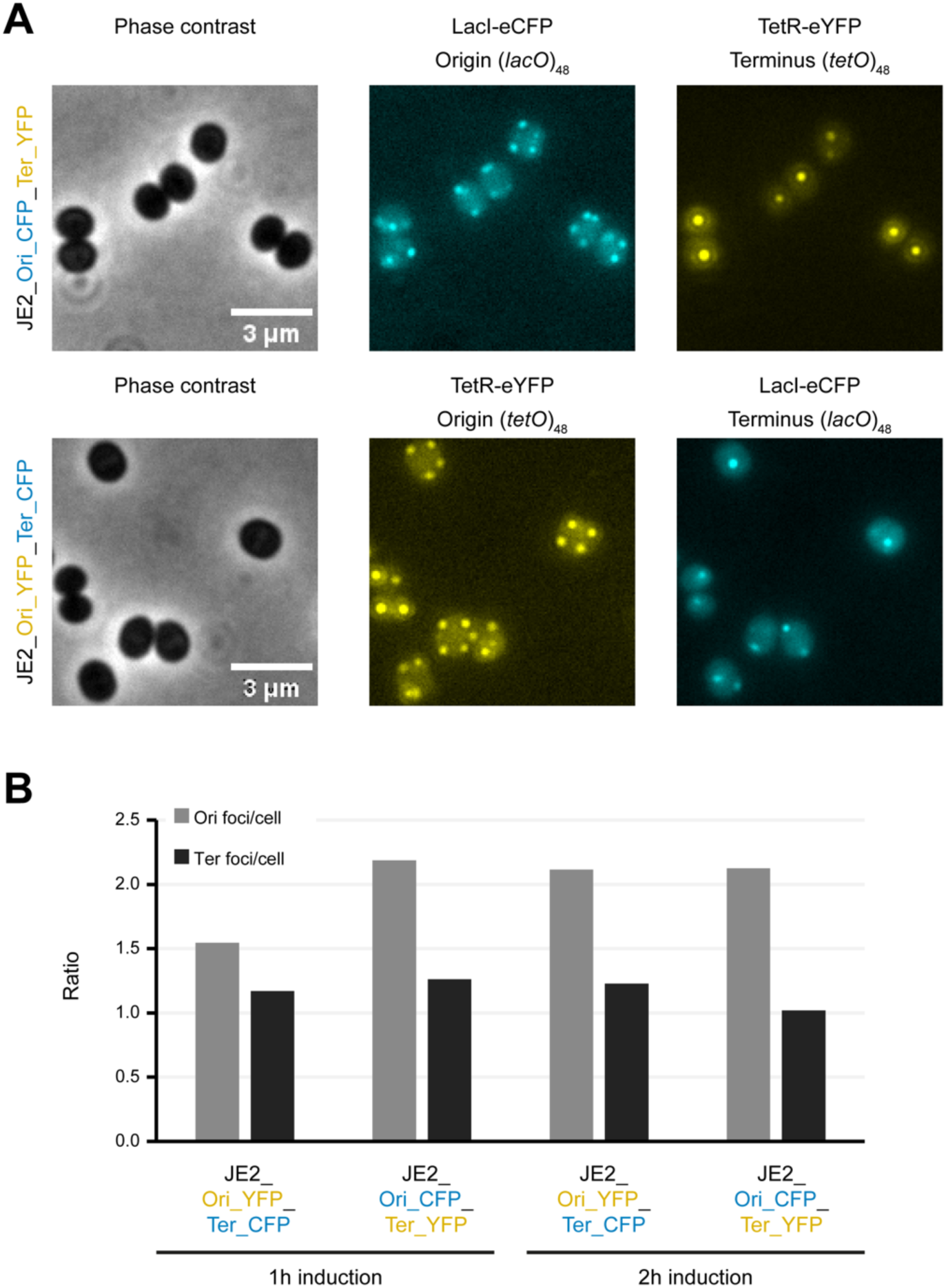
Dual labeling of origins and termini using the FROS system. **A** Phase contrast (left) and fluorescence (middle and right) microscopy images of cells expressing TerR-eYFP and LacI-eCFP in strain JE2_Ori_CFP_Ter_YFP (top) with a (*lacO*)_48_ array integrated near the origin and a (*tetO*)_48_ array near the terminus, and in strain JE2_Ori_YFP_Ter_CFP (bottom) with a (*lacO*)_48_ array integrated near the terminus and (*tetO*)_48_ array near the origin. **B** Bar chart showing the number of Ori/cell and Ter/cell for the strains showed in **A**, after 1 or 2 hours of induction with 1µM of CdCl_2_ (n>1200 per condition).

**Supplementary Figure 3.**
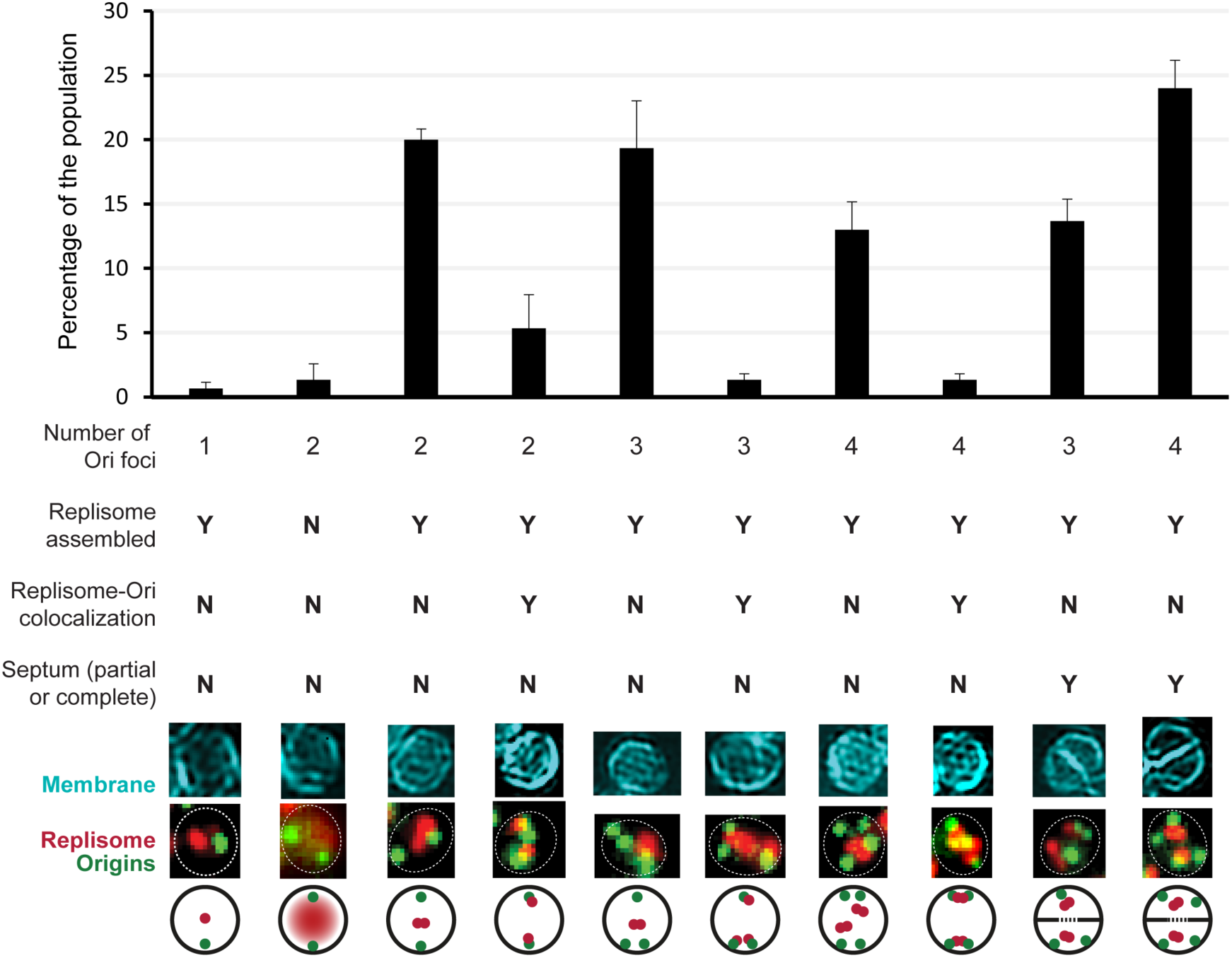
Origin number and co-localization with replisome during the *S. aureus* cell cycle. Bar chart showing the distribution of JE2_FROS^Ori^_DnaN-Halo cells manually classified according to the number of origin foci, replisome assembly, colocalization of DnaN-Halo with origins and presence of a visible septum (partial or complete). Data from three biological replicates, error bars indicate standard deviation, n=100. Below the chart are images of representative cells with origins labeled by TetR-mNG, DnaN-Halo labeled with JF549 and membrane labeled with CellBrite Fix 640 dye. The bottom row shows schematic representations of these cells to illustrate the classification.

**Supplementary Figure 4.**
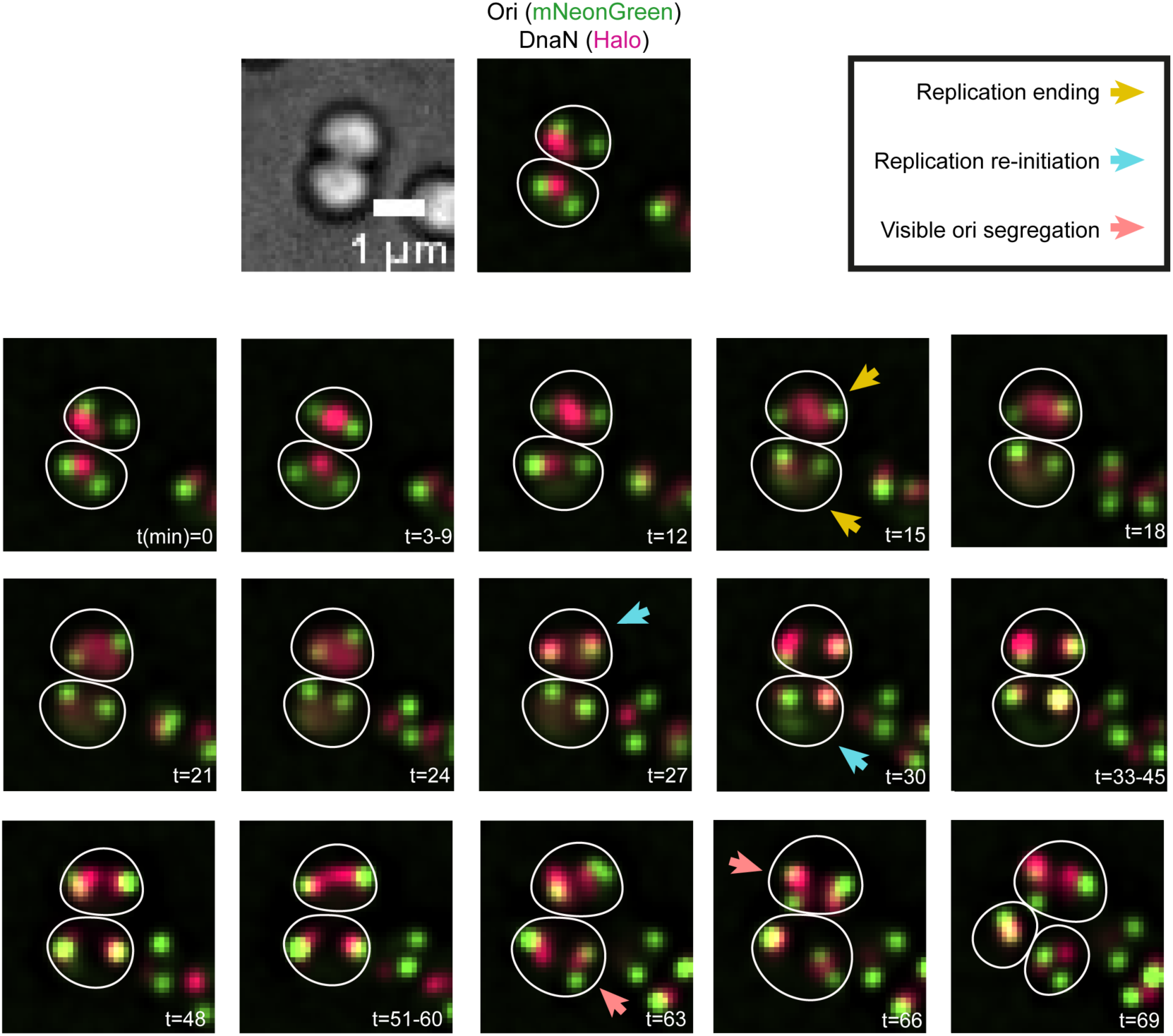
Representative time-lapse of the *S. aureus* chromosome cycle. Images from time-lapse microscopy of *S. aureus* cells of the strain JE2_FROS^Ori^_DnaN-Halo with origins labeled by TetR-mNeonGreen (green) and DnaN-Halo fusion labeled with JF549 (red). Frames were captured at 3 min intervals. For periods without relevant changes, only the initial frame is shown and the range is indicated in the lower right corner. Cell outlines are shown in white. Arrows mark key events: replication termination (yellow), re-initiation (blue) and visible origin segregation (red).

**Supplementary Figure 5.**
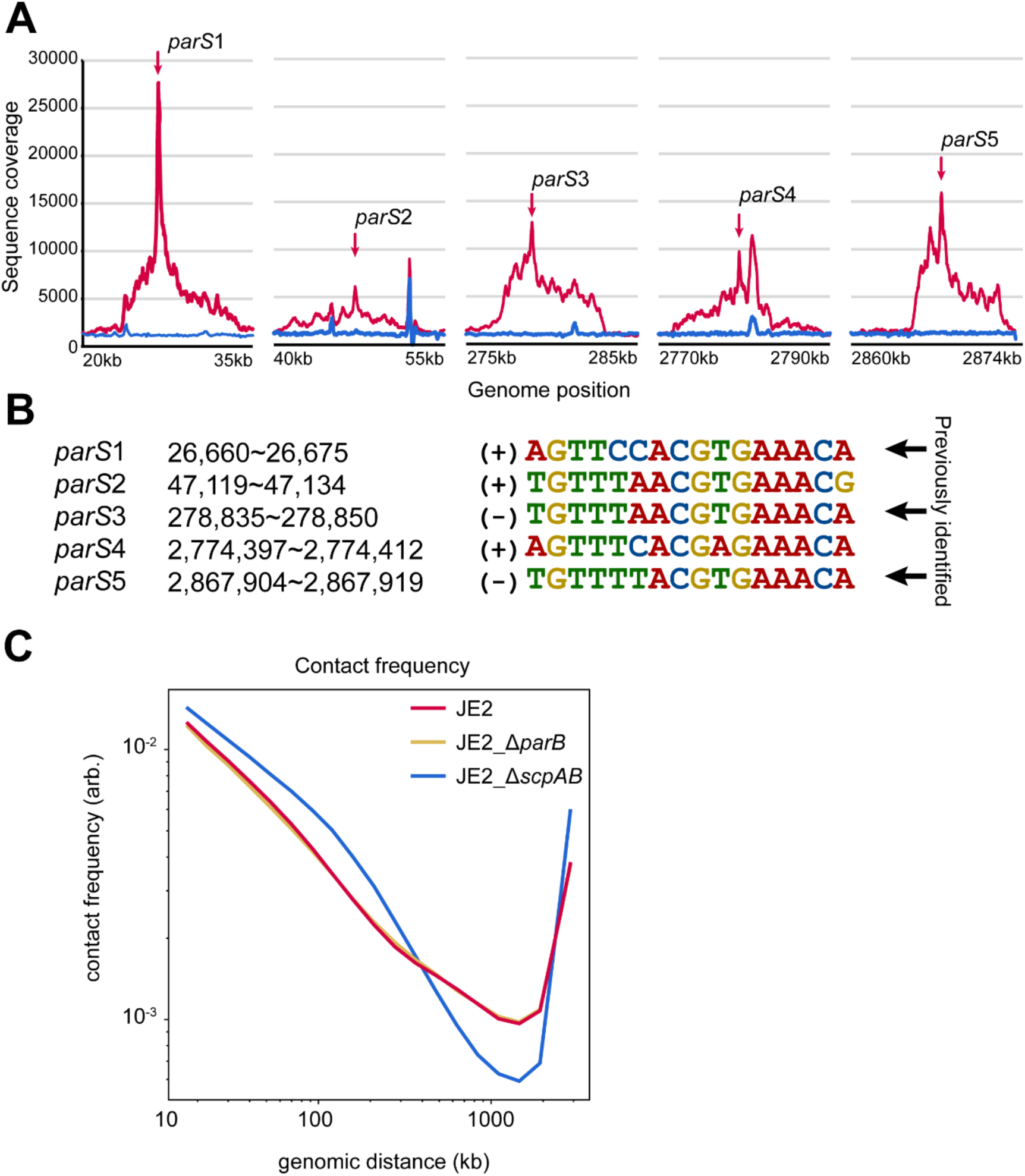
Identification of ParB binding sites and role of ParB and SMC-ScpAB complex in chromosome organization. **A** DNA regions enriched by anti-FLAG immunoprecipitation in JE2_ParB-3xFLAG (red) compared to JE2_3xFLAG-mNG control (blue). The x-axis indicates genomic position and the y-axis represents the number of reads for each position. Red arrows mark the position of *parS* sequences. **B** List of the five identified *parS* sites in *S. aureus*, including genomic position, sequence and whether the shown sequence is in the (+) or (-) strand of the chromosome. The sites *parS*1, *parS*3 and *parS*5 were previously described (*23*). **C** Contact probability curves for JE2 (red), JE2_Δ*parB* (yellow) and JE2_Δ*scpAB* (blue) strains. The curves represent the averaged probability of a contact (y-axis) between two loci separated by a given genomic distance indicated in the x-axis. Sequences separated by shorter distances (left side) have a higher probability of being in contact, and as the distance grows (moving towards the right side), the probability of contact decreases. Since the chromosome is circular, the curve rises again at its rightmost end.

**Supplementary Figure 6.**
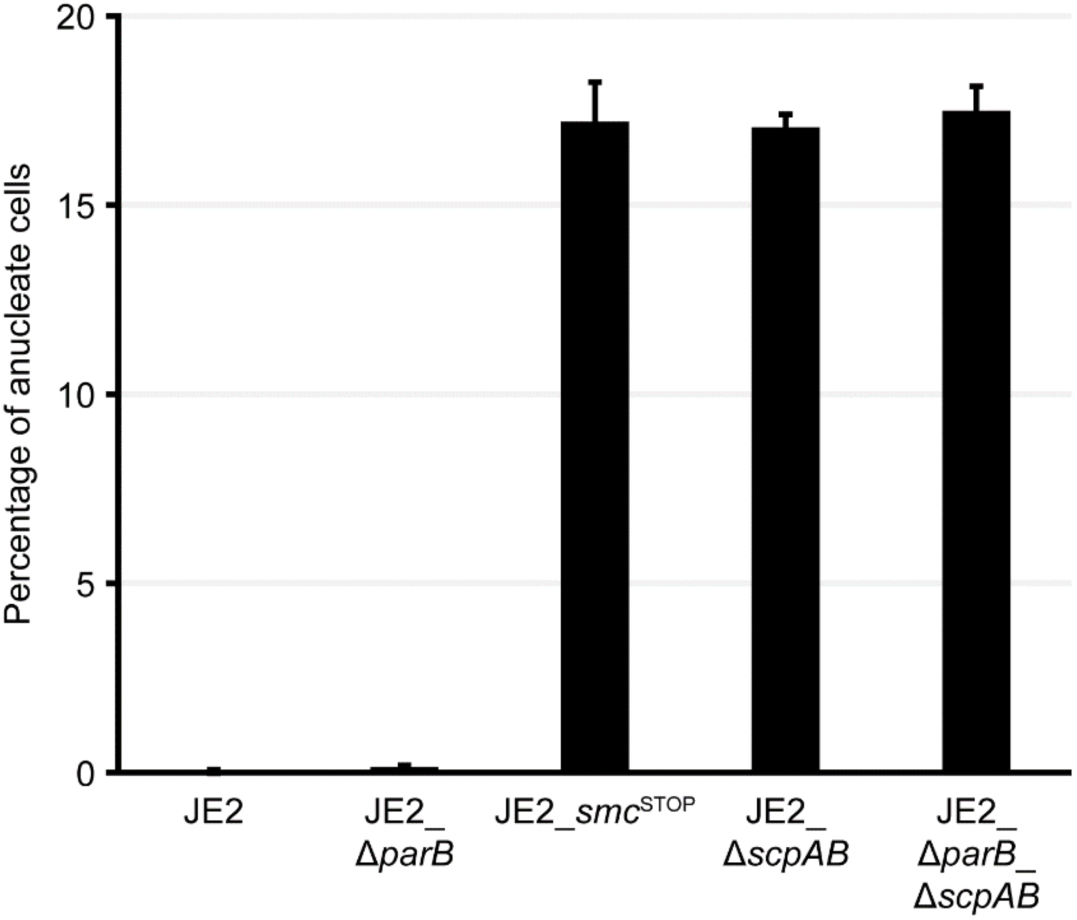
Quantification of anucleate cells in mutants lacking ParB or SMC-ScpAB. Bar graph showing the fraction of anucleate cells in the indicated strains. Data from three biological replicates, n>800 each. Error bars indicate standard deviation.

**Supplementary Figure 7.**
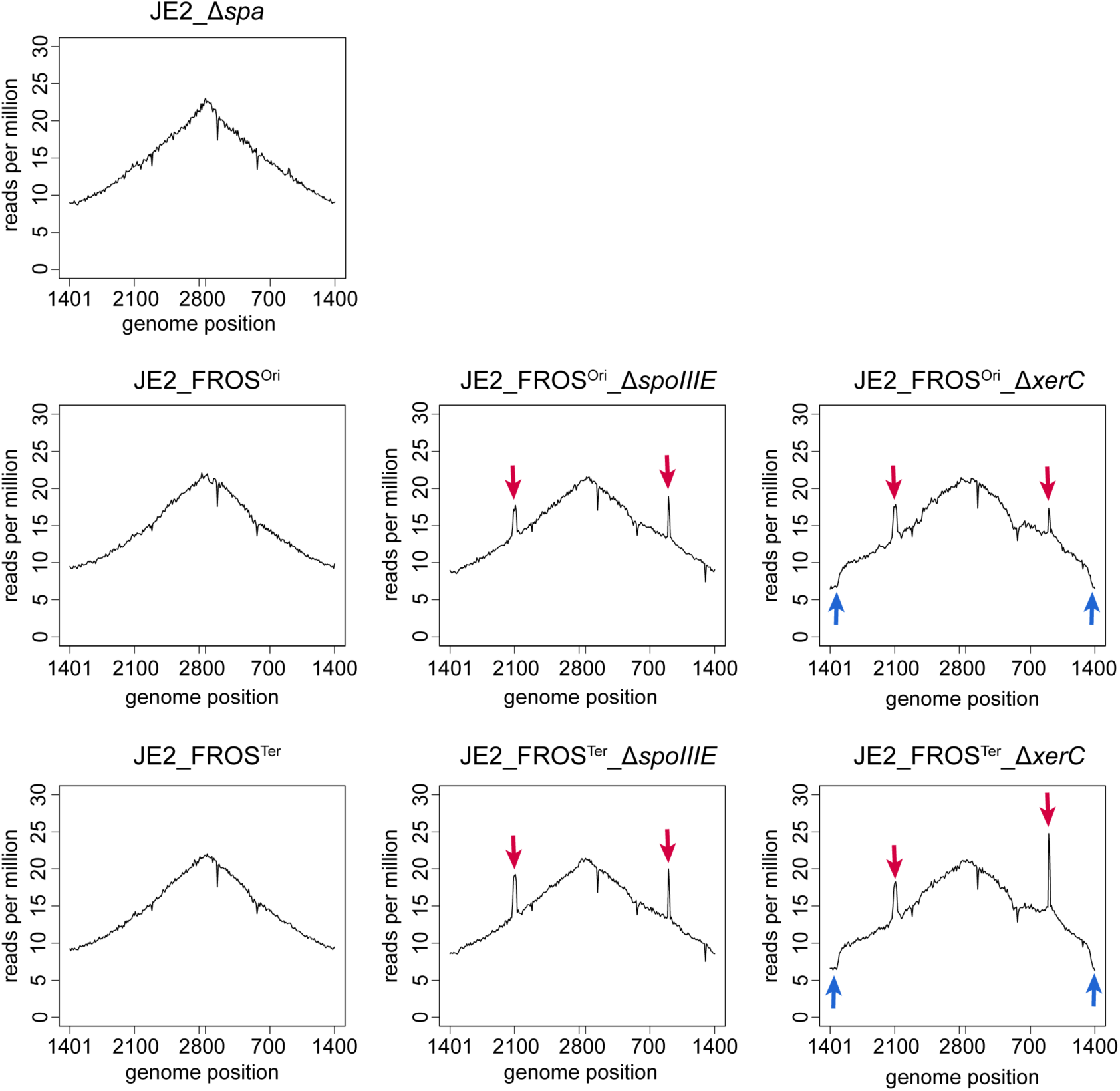
Genome-wide marker frequency analysis of *S. aureus* mutants lacking SpoIIIE or XerC. Marker frequency analysis based on whole genome sequencing of indicated strains. JE2_Δ*spa* was used as a reference as the FROS strains also lack the spa gene. The x axis represents the genomic position, with the origin at the center. Sequencing reads were normalized to the total number of reads for each sample (reads per million) and mapped in 10-kb bins. Red arrows mark peaks that are absent in the reference strain. Blue arrows indicate drops that are absent in the reference strain.

**Supplementary Table 1:**
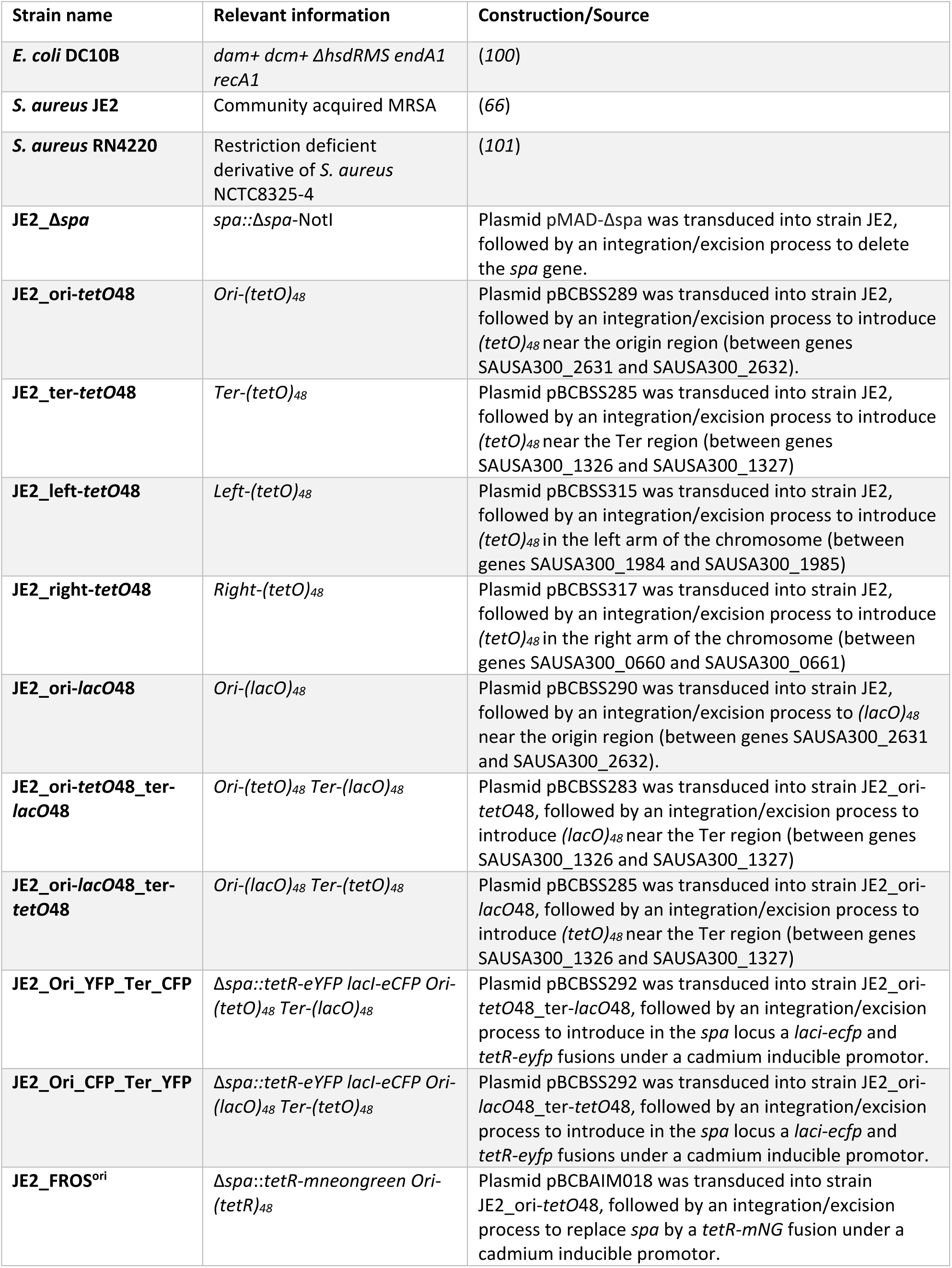
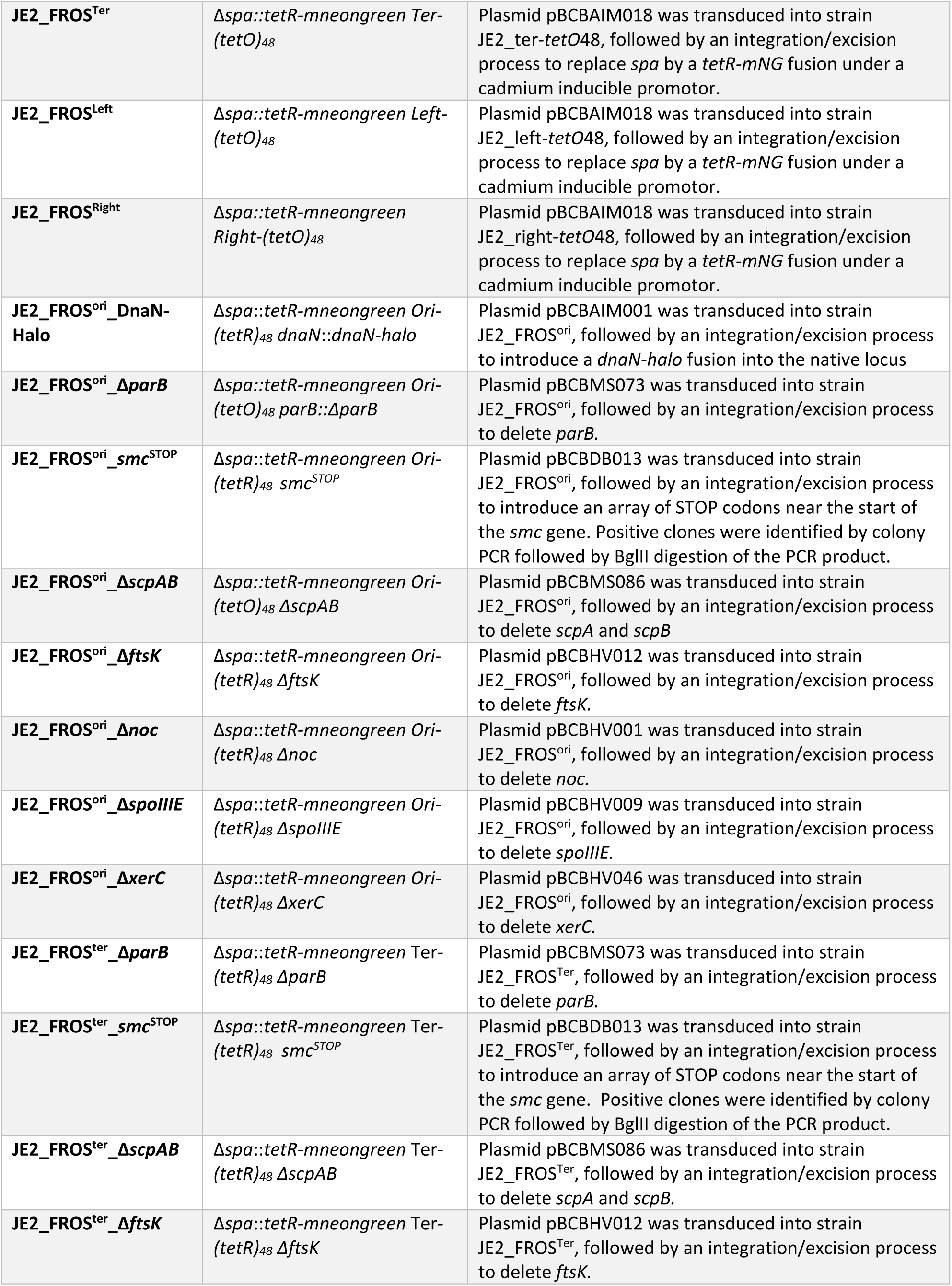
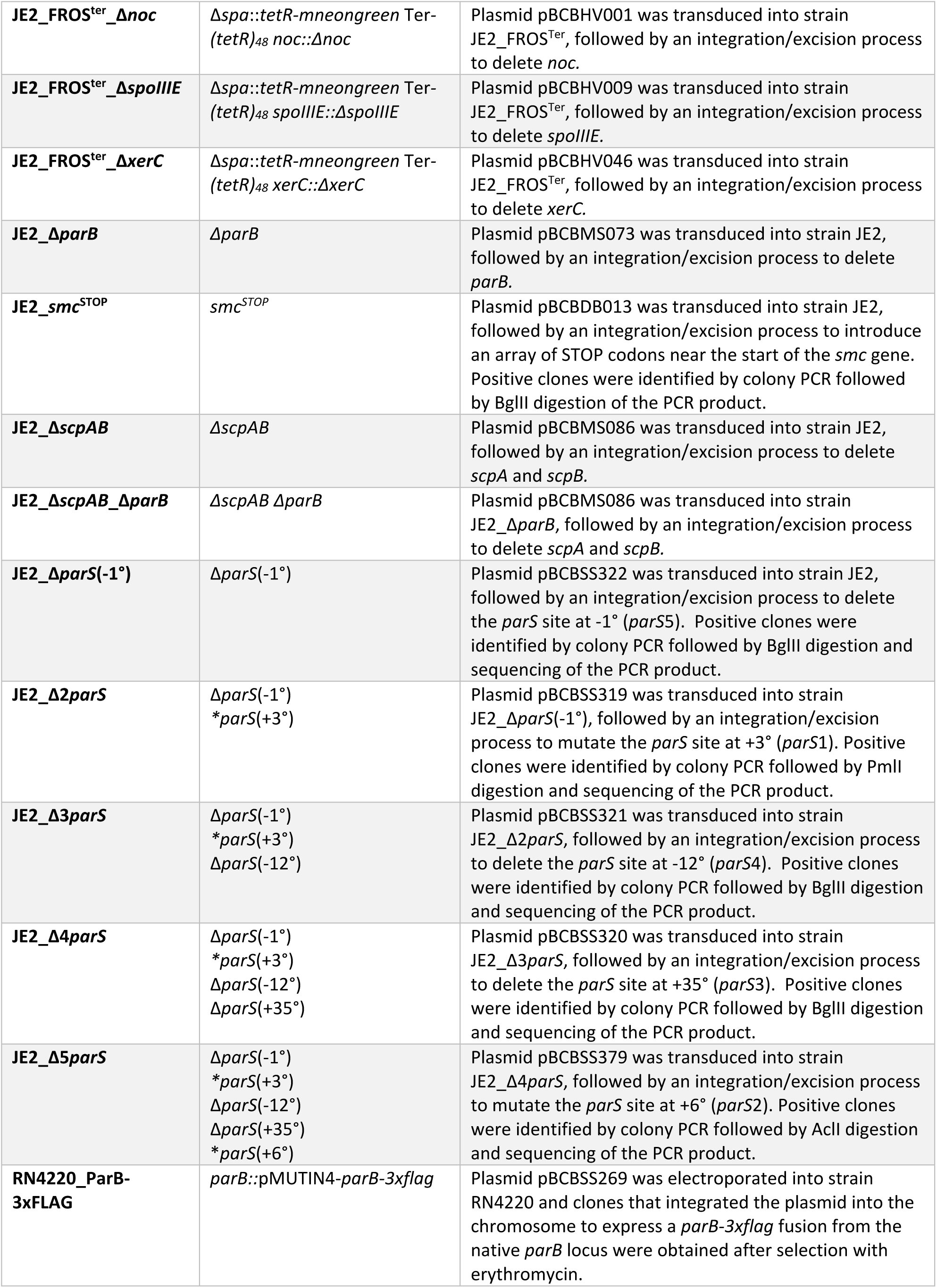
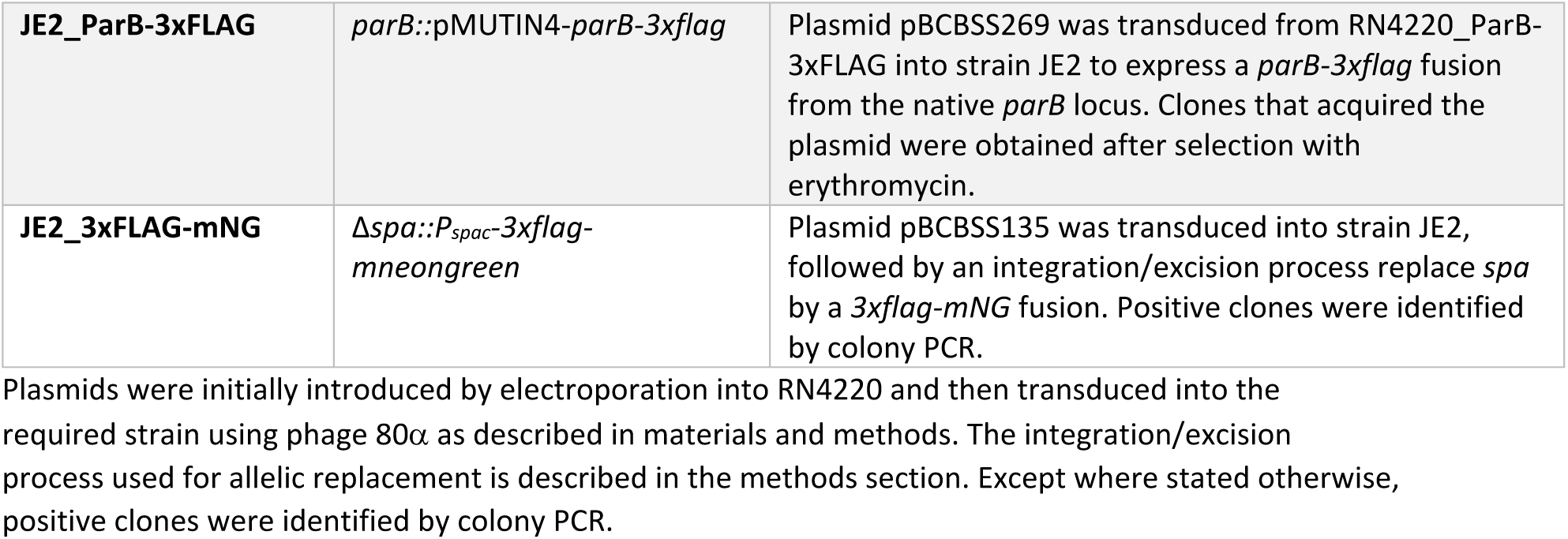
Strains used in the study.

**Supplementary Table 2:**
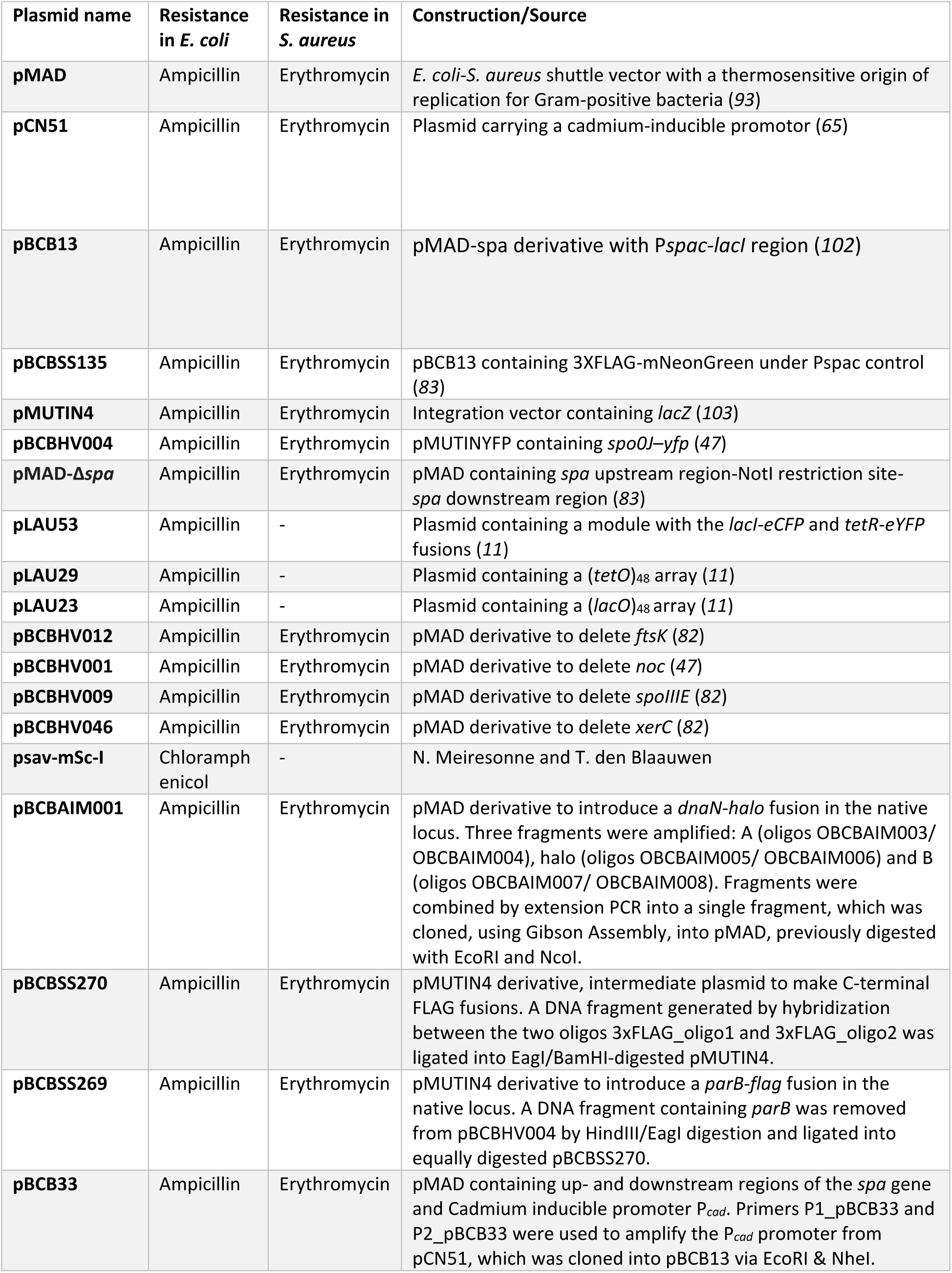
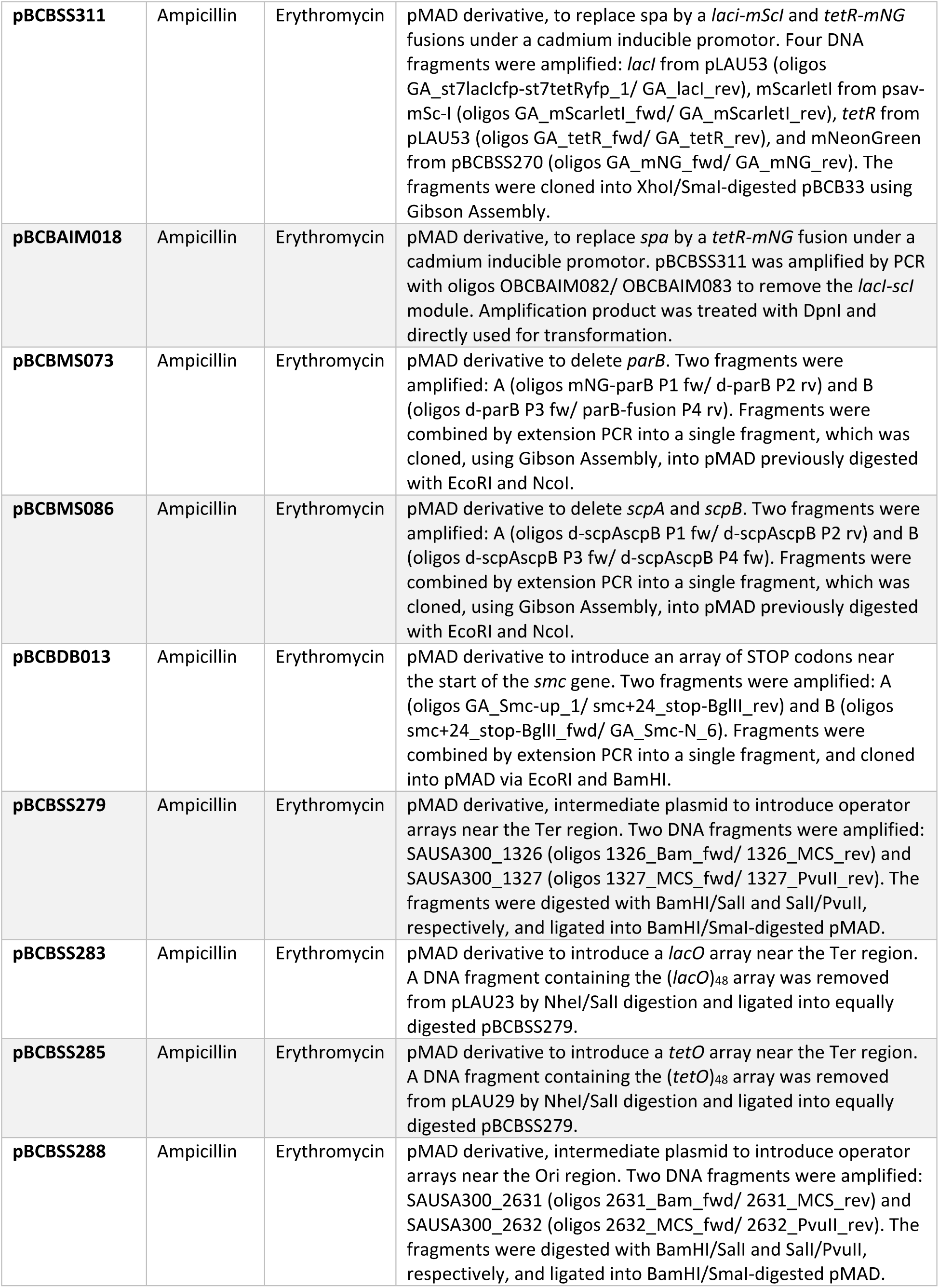
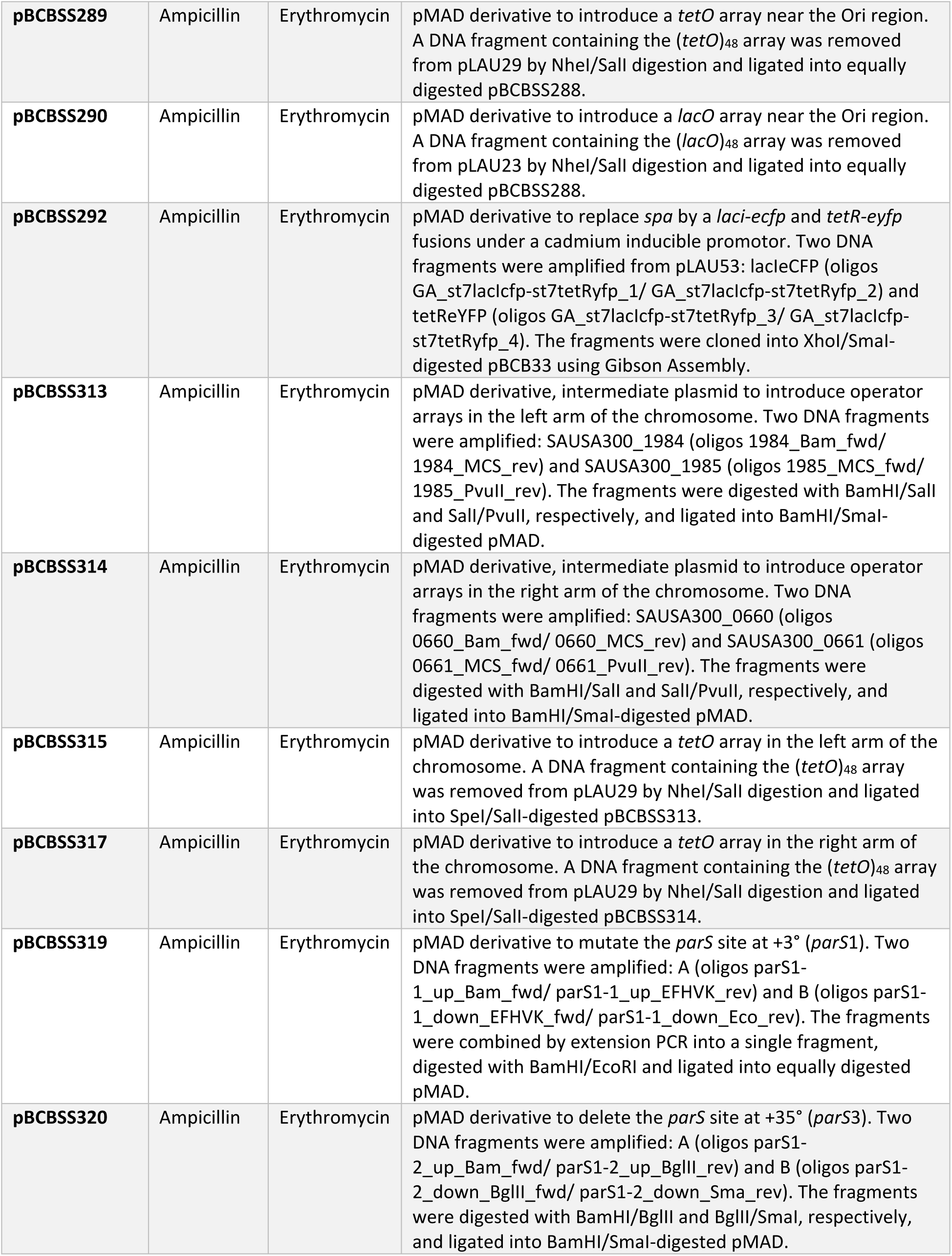
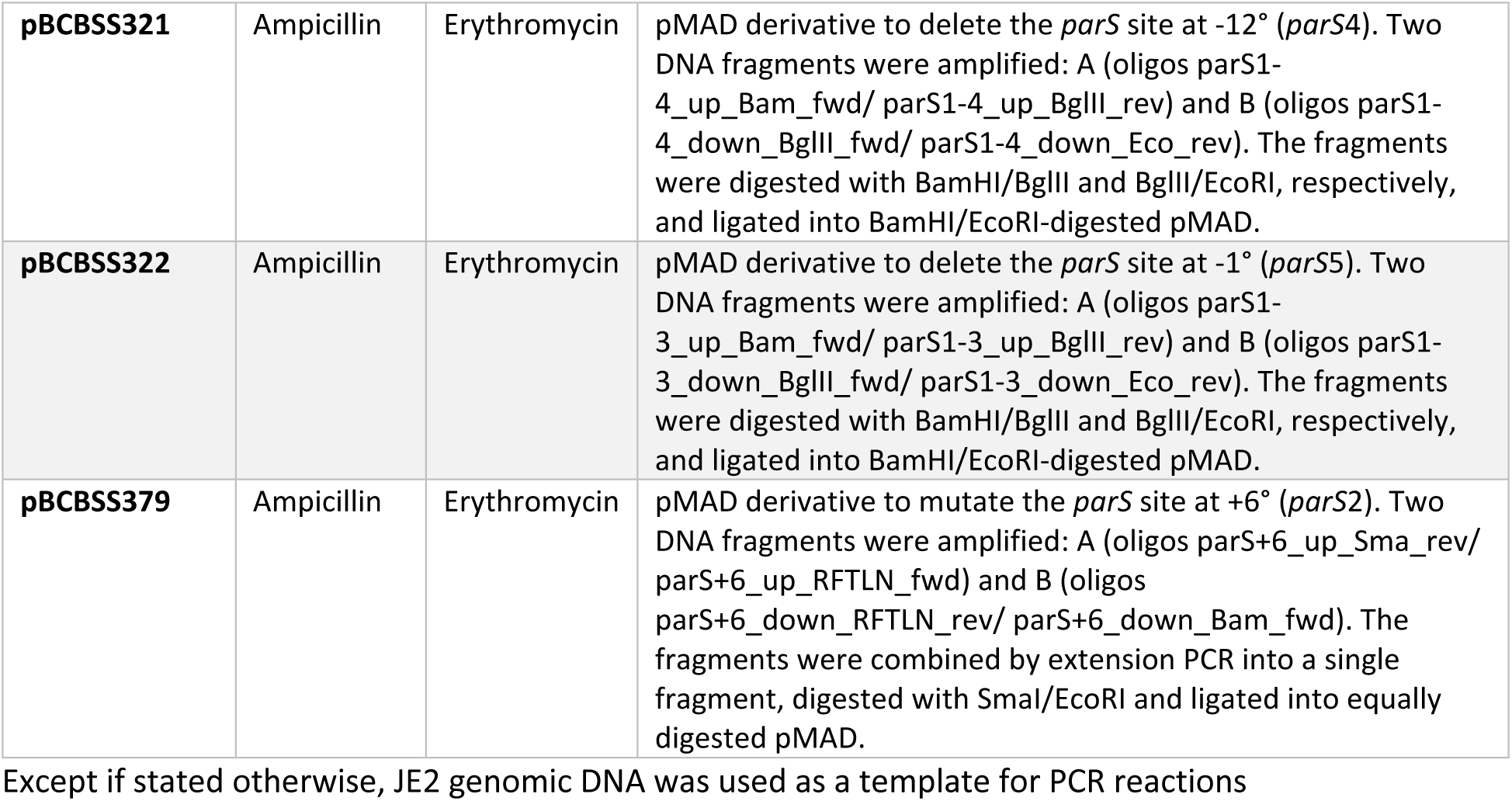
Plasmids used in the study.

**Supplementary Table 3:**
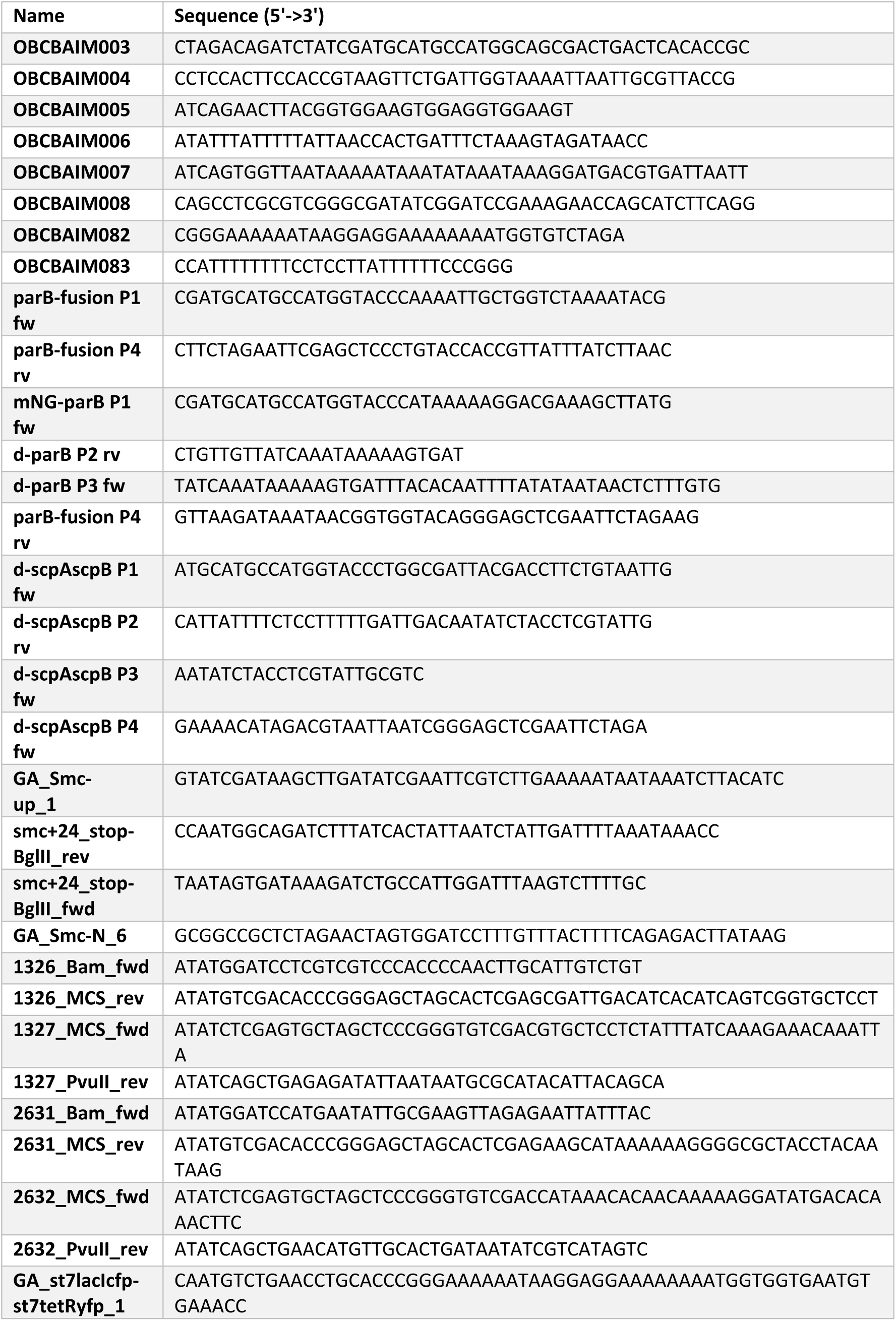
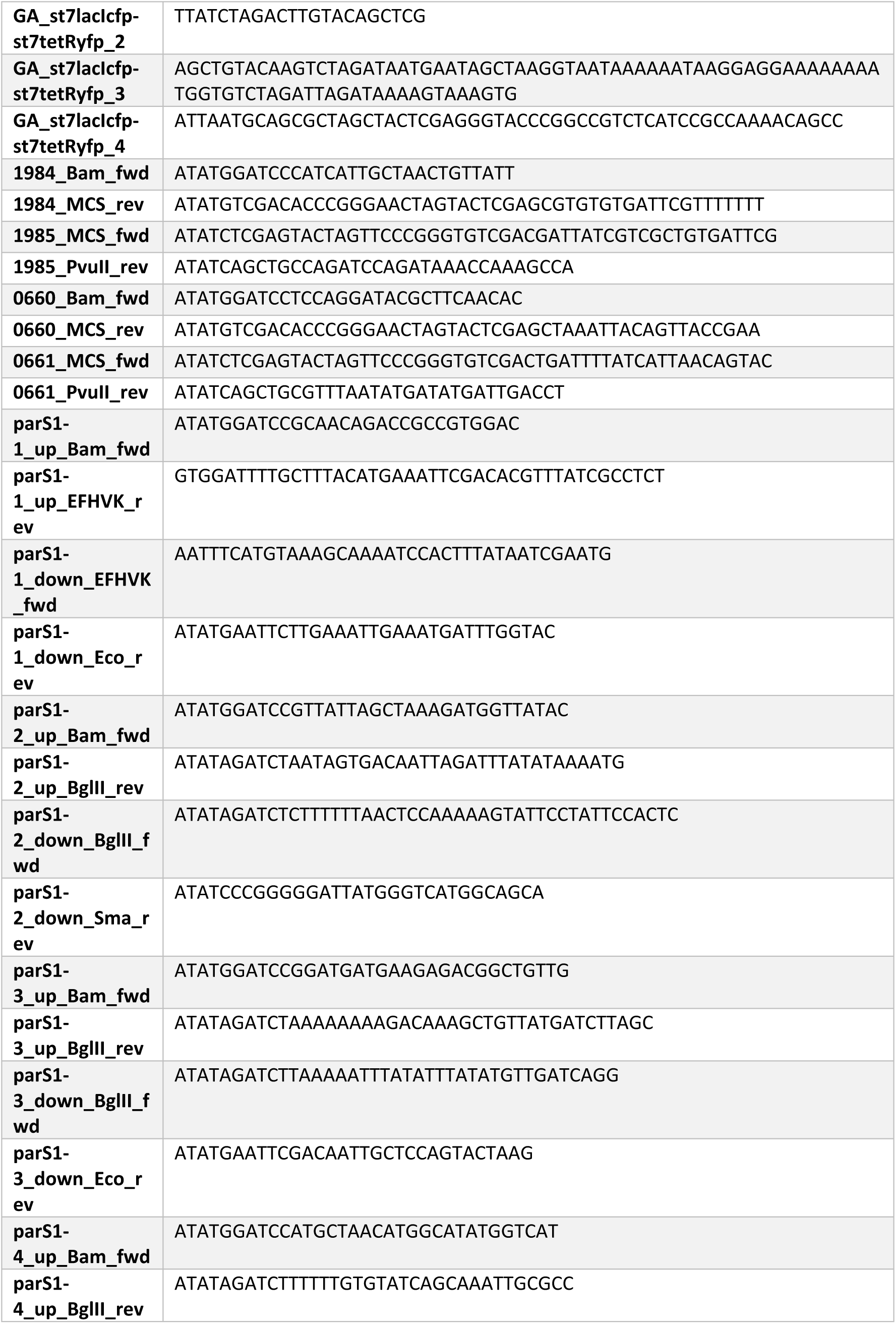
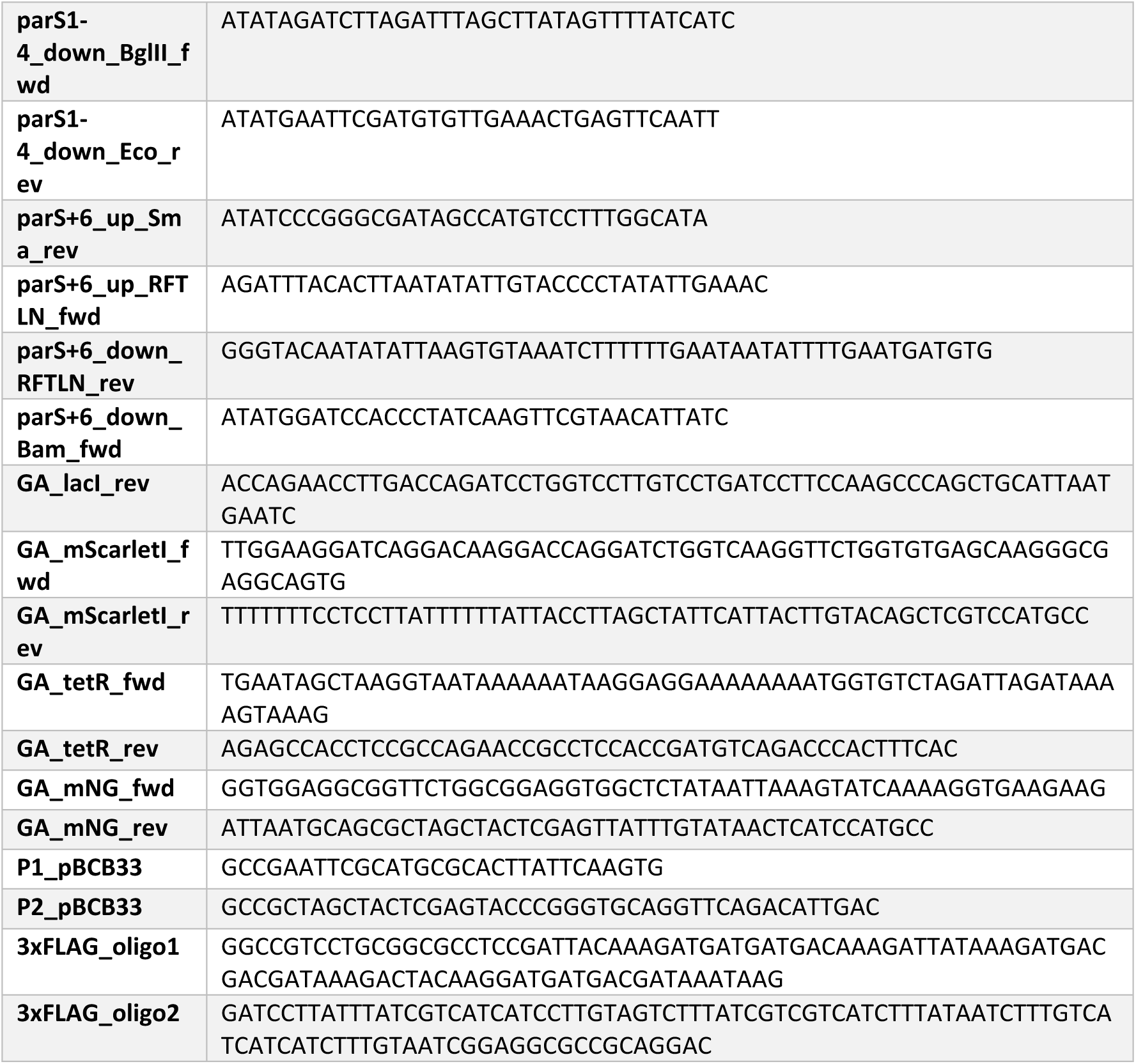
oligonucleotides used in the study.

## Notes

### Competing Interest Statement

The authors have declared no competing interest.

## Bibliography

1. C. Gogou, A. Japaridze, C. Dekker, Mechanisms for chromosome segregation in bacteria. Frontiers in Microbiology 12 (2021).

2. X. Wang, P. M. Llopis, D. Z. Rudner, Organization and segregation of bacterial chromosomes. Nature reviews. Genetics 14, 10.1038/nrg3375 (2013).

3. P. H. Viollier, M. Thanbichler, P. T. McGrath, L. West, M. Meewan, H. H. McAdams, L. Shapiro, Rapid and sequential movement of individual chromosomal loci to specific subcellular locations during bacterial DNA replication. Proceedings of the National Academy of Sciences 101, 9257–9262 (2004).

4. A. Harms, A. Treuner-Lange, D. Schumacher, L. Søgaard-Andersen, Tracking of chromosome and replisome dynamics in *Myxococcus xanthus* reveals a novel chromosome arrangement. PLOS Genetics 9, e1003802 (2013).

5. A. David, G. Demarre, L. Muresan, E. Paly, F.-X. Barre, C. Possoz, The two cis-acting sites, *parS1* and *oriC1*, contribute to the longitudinal organisation of *Vibrio cholerae* chromosome I. PLOS Genetics 10, e1004448 (2014).

6. X. Wang, P. Montero Llopis, D. Z. Rudner, *Bacillus subtilis* chromosome organization oscillates between two distinct patterns. Proceedings of the National Academy of Sciences 111, 12877–12882 (2014).

7. C. D. Webb, A. Teleman, S. Gordon, A. Straight, A. Belmont, D. C. Lin, A. D. Grossman, A. Wright, R. Losick, Bipolar localization of the replication origin regions of chromosomes in vegetative and sporulating cells of *Bacillus subtilis*. Cell 88, 667–674 (1997).

8. Böhm K, F. Meyer, A. Rhomberg, J. Kalinowski, C. Donovan, M. Bramkamp, Novel chromosome organization pattern in Actinomycetales-overlapping replication cycles combined with diploidy. mBio 8 (2017).

9. H. J. Nielsen, J. R. Ottesen, B. Youngren, S. J. Austin, F. G. Hansen, The *Escherichia coli* chromosome is organized with the left and right chromosome arms in separate cell halves. Molecular Microbiology 62, 331–338 (2006).

10. X. Wang, X. Liu, C. Possoz, D. J. Sherratt, The two *Escherichia coli* chromosome arms locate to separate cell halves. Genes & Development 20, 1727–1731 (2006).

11. I. F. Lau, S. R. Filipe, B. Søballe, O.-A. Økstad, F.-X. Barre, D. J. Sherratt, Spatial and temporal organization of replicating *Escherichia coli* chromosomes. Molecular Microbiology 49, 731–743 (2003).

12. K. Gras, D. Fange, J. Elf, The *Escherichia coli* chromosome moves to the replisome. Nature Communications 15, 6018 (2024).

13. R. van Raaphorst, M. Kjos, J.-W. Veening, Chromosome segregation drives division site selection in *Streptococcus pneumoniae*. Proceedings of the National Academy of Sciences 114, E5959–E5968 (2017).

14. I. Santi, J. D. McKinney, Chromosome organization and replisome dynamics in *Mycobacterium smegmatis*. mBio 6, e01999–14 (2015).

15. H. Yoshikawa, A. O’Sullivan, N. Sueoka, Sequential replication of the *Bacillus subtilis* chromosome, III. regulation of initiation. Proceedings of the National Academy of Sciences 52, 973–980 (1964).

16. S. Cooper, C. E. Helmstetter, Chromosome replication and the division cycle of *Escherichia coli* B/r. Journal of Molecular Biology 31, 519–540 (1968).

17. D. Trojanowski, J. Hołówka, K. Ginda, D. Jakimowicz, J. Zakrzewska-Czerwińska, Multifork chromosome replication in slow-growing bacteria. Scientific Reports 7, 43836 (2017).

18. D. C. Lin, A. D. Grossman, Identification and characterization of a bacterial chromosome partitioning site. Cell 92, 675–685 (1998).

19. E. Toro, S.-H. Hong, H. H. McAdams, L. Shapiro, *Caulobacter* requires a dedicated mechanism to initiate chromosome segregation. Proceedings of the National Academy of Sciences 105, 15435–15440 (2008).

20. R. Kadoya, J. H. Baek, A. Sarker, D. K. Chattoraj, Participation of chromosome segregation protein ParAI of *Vibrio cholerae* in chromosome replication. Journal of Bacteriology 193, 1504–1514 (2011).

21. A. A. Iniesta, ParABS system in chromosome partitioning in the bacterium *Myxococcus xanthus*. PLOS ONE 9, e86897 (2014).

22. A. S. B. Jalal, T. B. K. Le, Bacterial chromosome segregation by the ParABS system. Open Biology 10, 200097 (2020).

23. J. Livny, Y. Yamaichi, M. K. Waldor, Distribution of centromere-like *parS* sites in Bacteria: insights from comparative genomics. Journal of Bacteriology 189, 8693–8703 (2007).

24. S. Gruber, J. Errington, Recruitment of condensin to replication origin regions by ParB/SpoOJ promotes chromosome segregation in *B. subtilis*. Cell 137, 685–696 (2009).

25. N. L. Sullivan, K. A. Marquis, D. Z. Rudner, Recruitment of SMC by ParB-*parS* organizes the origin region and promotes efficient chromosome segregation. Cell 137, 697–707 (2009).

26. N. T. Tran, M. T. Laub, T. B. K. Le, SMC progressively aligns chromosomal arms in *Caulobacter crescentus* but is antagonized by convergent transcription. Cell Reports 20, 2057–2071 (2017).

27. S. Gruber, J.-W. Veening, J. Bach, M. Blettinger, M. Bramkamp, J. Errington, Interlinked sister chromosomes arise in the absence of condensin during fast replication in *B. subtilis*. Current Biology 24, 293 (2014).

28. J. Mäkelä, D. Sherratt, SMC complexes organize the bacterial chromosome by lengthwise compaction. Current Genetics 66, 895–899 (2020).

29. X. Wang, H. B. Brandão, T. B. K. Le, M. T. Laub, D. Z. Rudner, *Bacillus subtilis* SMC complexes juxtapose chromosome arms as they travel from origin to terminus. Science 355, 524–527 (2017).

30. X. Wang, O. W. Tang, E. P. Riley, D. Z. Rudner, The SMC condensin complex is required for origin segregation in *Bacillus subtilis*. Current Biology 24, 287–292 (2014).

31. D. Krepel, R. R. Cheng, M. Di Pierro, J. N. Onuchic, Deciphering the structure of the condensin protein complex. Proceedings of the National Academy of Sciences 115, 11911–11916 (2018).

32. L. Wilhelm, F. Bürmann, A. Minnen, H.-C. Shin, C. P. Toseland, B.-H. Oh, S. Gruber, SMC condensin entraps chromosomal DNA by an ATP hydrolysis dependent loading mechanism in *Bacillus subtilis*. eLife 4, e06659 (2015).

33. K. Böhm, G. Giacomelli, A. Schmidt, A. Imhof, R. Koszul, M. Marbouty, M. Bramkamp, Chromosome organization by a conserved condensin-ParB system in the actinobacterium *Corynebacterium glutamicum*. Nature Communications 11, 1485 (2020).

34. V. S. Lioy, I. Junier, V. Lagage, I. Vallet, F. Boccard, Distinct activities of bacterial condensins for chromosome management in *Pseudomonas aeruginosa*. Cell Reports 33, 108344 (2020).

35. M. A. Schwartz, L. Shapiro, An SMC ATPase mutant disrupts chromosome segregation in *Caulobacter*. Molecular Microbiology 82, 1359–1374 (2011).

36. D. Anand, D. Schumacher, L. Søgaard-Andersen, SMC and the bactofilin/PadC scaffold have distinct yet redundant functions in chromosome segregation and organization in *Myxococcus xanthus*. Molecular Microbiology 114, 839–856 (2020).

37. C. Donovan, A. Schwaiger, R. Krämer, M. Bramkamp, Subcellular localization and characterization of the ParAB system from *Corynebacterium glutamicum*. Journal of Bacteriology 192, 3441–3451 (2010).

38. C. Güthlein, R. M. Wanner, P. Sander, E. C. Böttger, B. Springer, A mycobacterial *smc* null mutant is proficient in DNA repair and long-term survival. Journal of Bacteriology 190, 452–456 (2008).

39. D. Jakimowicz, A. Brzostek, A. Rumijowska-Galewicz, P. Żydek, A. Dołzbłasz, A. Smulczyk-Krawczyszyn, T. Zimniak, Ł. Wojtasz, A. Zawilak-Pawlik, A. Kois, J. Dziadek, J. Zakrzewska-Czerwińska, Characterization of the mycobacterial chromosome segregation protein ParB and identification of its target in *Mycobacterium smegmatis*. Microbiology 153, 4050– 4060 (2007).

40. P. Jecz, A. A. Bartosik, K. Glabski, G. Jagura-Burdzy, A single *parS* sequence from the cluster of four sites closest to *oriC* is necessary and sufficient for proper chromosome segregation in *Pseudomonas aeruginosa*. PloS One 10, e0120867 (2015).

41. R. B. Jensen, L. Shapiro, The *Caulobacter crescentus smc* gene is required for cell cycle progression and chromosome segregation. Proceedings of the National Academy of Sciences 96, 10661–10666 (1999).

42. A. Jung, A. Raßbach, R. L. Pulpetta, M. C. F. van Teeseling, K. Heinrich, P. Sobetzko, J. Serrania, A. Becker, M. Thanbichler, Two-step chromosome segregation in the stalked budding bacterium *Hyphomonas neptunium*. Nature Communications 10, 3290 (2019).

43. A. Minnen, L. Attaiech, M. Thon, S. Gruber, J.-W. Veening, SMC is recruited to *oriC* by ParB and promotes chromosome segregation in *Streptococcus pneumoniae*. Molecular Microbiology 81, 676–688 (2011).

44. S. Jun, A. Wright, Entropy as the driver of chromosome segregation. Nature reviews. Microbiology 8, 600–607 (2010).

45. T. G. Bernhardt, P. A. J. de Boer, SlmA, a nucleoid-associated, FtsZ binding protein required for blocking septal ring assembly over chromosomes in *E. coli*. Molecular Cell 18, 555–564 (2005).

46. M. Thanbichler, L. Shapiro, MipZ, a spatial regulator coordinating chromosome segregation with cell division in *Caulobacter*. Cell 126, 147–162 (2006).

47. H. Veiga, A. M. Jorge, M. G. Pinho, Absence of nucleoid occlusion effector Noc impairs formation of orthogonal FtsZ rings during *Staphylococcus aureus* cell division. Molecular Microbiology 80, 1366–1380 (2011).

48. L. J. Wu, S. Ishikawa, Y. Kawai, T. Oshima, N. Ogasawara, J. Errington, Noc protein binds to specific DNA sequences to coordinate cell division with chromosome segregation. The EMBO Journal 28, 1940–1952 (2009).

49. L. J. Wu, J. Errington, Coordination of cell division and chromosome segregation by a nucleoid occlusion protein in *Bacillus subtilis*. Cell 117, 915–925 (2004).

50. K. S. Ramamurthi, S. Lecuyer, H. A. Stone, R. Losick, Geometric cue for protein localization in a bacterium. Science 323, 1354–1357 (2009).

51. L. Shapiro, H. H. McAdams, R. Losick, Why and how bacteria localize proteins. Science 326, 1225–1228 (2009).

52. K. S. Ramamurthi, R. Losick, Negative membrane curvature as a cue for subcellular localization of a bacterial protein. Proceedings of the National Academy of Sciences 106, 13541–13545 (2009).

53. A. Varma, K. C. Huang, K. D. Young, The Min system as a general cell geometry detection mechanism: branch lengths in Y-shaped *Escherichia coli* cells affect Min oscillation patterns and division dynamics. Journal of Bacteriology 190, 2106–2117 (2008).

54. P. A. de Boer, R. E. Crossley, L. I. Rothfield, A division inhibitor and a topological specificity factor coded for by the minicell locus determine proper placement of the division septum in *E. coli*. Cell 56, 641–649 (1989).

55. S. Hussain, C. N. Wivagg, P. Szwedziak, F. Wong, K. Schaefer, T. Izoré, L. D. Renner, M. J. Holmes, Y. Sun, A. W. Bisson-Filho, S. Walker, A. Amir, J. Löwe, E. C. Garner, MreB filaments align along greatest principal membrane curvature to orient cell wall synthesis. eLife 7, e32471 (2018).

56. J. A. Taylor, B. P. Bratton, S. R. Sichel, K. M. Blair, H. M. Jacobs, K. E. DeMeester, E. Kuru, J. Gray, J. Biboy, M. S. VanNieuwenhze, W. Vollmer, C. L. Grimes, J. W. Shaevitz, N. R. Salama, Distinct cytoskeletal proteins define zones of enhanced cell wall synthesis in *Helicobacter pylori*. eLife 9, e52482 (2020).

57. T. S. Ursell, J. Nguyen, R. D. Monds, A. Colavin, G. Billings, N. Ouzounov, Z. Gitai, J. W. Shaevitz, K. C. Huang, Rod-like bacterial shape is maintained by feedback between cell curvature and cytoskeletal localization. Proceedings of the National Academy of Sciences 111, E1025–1034 (2014).

58. A. S. Lee, H. de Lencastre, J. Garau, J. Kluytmans, S. Malhotra-Kumar, A. Peschel, S. Harbarth, Methicillin-resistant *Staphylococcus aureus*. Nature Reviews Disease Primers 4, 1–23 (2018).

59. Antimicrobial Resistance Collaborators, Global burden of bacterial antimicrobial resistance in 2019: a systematic analysis. Lancet 399, 629–655 (2022).

60. J. M. Monteiro, P. B. Fernandes, F. Vaz, A. R. Pereira, A. C. Tavares, M. T. Ferreira, P. M. Pereira, H. Veiga, E. Kuru, M. S. VanNieuwenhze, Y. V. Brun, S. R. Filipe, M. G. Pinho, Cell shape dynamics during the staphylococcal cell cycle. Nature Communications 6, 8055 (2015).

61. M. G. Pinho, S. J. Foster, Cell growth and division of *Staphylococcus aureus*. Annual Review of Microbiology 78, 293–310 (2024).

62. B. M. Saraiva, M. Sorg, A. R. Pereira, M. J. Ferreira, L. C. Caulat, N. T. Reichmann, M. G. Pinho, Reassessment of the distinctive geometry of *Staphylococcus aureus* cell division. Nature Communications 11, 4097 (2020).

63. H. Chan, B. Söderström, U. Skoglund, Spo0J and SMC are required for normal chromosome segregation in *Staphylococcus aureus*. MicrobiologyOpen 9, e999 (2020).

64. M. G. Pinho, J. Errington, A divIVA null mutant of *Staphylococcus aureus* undergoes normal cell division. FEMS Microbiology Letters 240, 145–149 (2004).

65. E. Charpentier, A. I. Anton, P. Barry, B. Alfonso, Y. Fang, R. P. Novick, Novel cassette-based shuttle vector system for Gram-positive bacteria. Applied and Environmental Microbiology 70, 6076–6085 (2004).

66. P. D. Fey, J. L. Endres, V. K. Yajjala, T. J. Widhelm, R. J. Boissy, J. L. Bose, K. W. Bayles, A genetic resource for rapid and comprehensive phenotype screening of nonessential *Staphylococcus aureus* genes. mBio 4, 10.1128/mbio.00537-12 (2013).

67. N. C. Shaner, G. G. Lambert, A. Chammas, Y. Ni, P. J. Cranfill, M. A. Baird, B. R. Sell, J. R. Allen, R. N. Day, M. Israelsson, M. W. Davidson, J. Wang, A bright monomeric green fluorescent protein derived from *Branchiostoma lanceolatum*. Nature Methods 10, 407– 409 (2013).

68. G. V. Los, L. P. Encell, M. G. McDougall, D. D. Hartzell, N. Karassina, C. Zimprich, M. G. Wood, R. Learish, R. F. Ohana, M. Urh, D. Simpson, J. Mendez, K. Zimmerman, P. Otto, G. Vidugiris, J. Zhu, A. Darzins, D. H. Klaubert, R. F. Bulleit, K. V. Wood, HaloTag: a novel protein labeling technology for cell imaging and protein analysis. ACS Chemical Biology 3, 373–382 (2008).

69. P. J. Chen, A. B. McMullin, B. J. Visser, Q. Mei, S. M. Rosenberg, D. Bates, Interdependent progression of bidirectional sister replisomes in *E. coli*. eLife 12, e82241 (2023).

70. B. M. Saraiva, L. Krippahl, S. R. Filipe, R. Henriques, M. G. Pinho, eHooke: A tool for automated image analysis of spherical bacteria based on cell cycle progression. Biological Imaging 1, e3 (2021).

71. J.-Y. Tinevez, N. Perry, J. Schindelin, G. M. Hoopes, G. D. Reynolds, E. Laplantine, S. Y. Bednarek, S. L. Shorte, K. W. Eliceiri, TrackMate: An open and extensible platform for single-particle tracking. Methods 115, 80–90 (2017).

72. T. B. K. Le, M. V. Imakaev, L. A. Mirny, M. T. Laub, High-resolution mapping of the spatial organization of a bacterial chromosome. Science 342, 731–734 (2013).

73. X. Karaboja, Z. Ren, H. B. Brandão, P. Paul, D. Z. Rudner, X. Wang, XerD unloads bacterial SMC complexes at the replication terminus. Molecular Cell 81, 756–766.e8 (2021).

74. J. Soppa, K. Kobayashi, M.-F. Noirot-Gros, D. Oesterhelt, S. D. Ehrlich, E. Dervyn, N. Ogasawara, S. Moriya, Discovery of two novel families of proteins that are proposed to interact with prokaryotic SMC proteins, and characterization of the *Bacillus subtilis* family members ScpA and ScpB. Molecular Microbiology 45, 59–71 (2002).

75. T. Pang, X. Wang, H. C. Lim, T. G. Bernhardt, D. Z. Rudner, The nucleoid occlusion factor Noc controls DNA replication initiation in *Staphylococcus aureus*. PLOS Genetics 13, e1006908 (2017).

76. R. R. Chaudhuri, A. G. Allen, P. J. Owen, G. Shalom, K. Stone, M. Harrison, T. A. Burgis, M. Lockyer, J. Garcia-Lara, S. J. Foster, S. J. Pleasance, S. E. Peters, D. J. Maskell, I. G. Charles, Comprehensive identification of essential *Staphylococcus aureus* genes using Transposon-Mediated Differential Hybridisation (TMDH). BMC genomics 10, 291 (2009).

77. A. L. Bottomley, A. T. F. Liew, K. D. Kusuma, E. Peterson, L. Seidel, S. J. Foster, E. J. Harry, Coordination of chromosome segregation and cell division in *Staphylococcus aureus*. Frontiers in Microbiology 8, 1575 (2017).

78. W. Yu, S. Herbert, P. L. Graumann, F. Götz, Contribution of SMC (Structural Maintenance of Chromosomes) and SpoIIIE to chromosome segregation in Staphylococci. Journal of Bacteriology 192, 4067–4073 (2010).

79. F. Ramos-León, B. R. Anjuwon-Foster, V. Anantharaman, T. B. Updegrove, C. N. Ferreira, A. M. Ibrahim, C.-H. Tai, M. J. Kruhlak, D. M. Missiakas, J. L. Camberg, L. Aravind, K. S. Ramamurthi, PcdA promotes orthogonal division plane selection in *Staphylococcus aureus*. Nature Microbiology 9, 2997–3012 (2024).

80. S. Bigot, V. Sivanathan, C. Possoz, F.-X. Barre, F. Cornet, FtsK, a literate chromosome segregation machine. Molecular Microbiology 64, 1434–1441 (2007).

81. L. Aussel, F.-X. Barre, M. Aroyo, A. Stasiak, A. Z. Stasiak, D. Sherratt, FtsK Is a DNA motor protein that activates chromosome dimer resolution by switching the catalytic state of the XerC and XerD recombinases. Cell 108, 195–205 (2002).

82. H. Veiga, M. G. Pinho, *Staphylococcus aureus* requires at least one FtsK/SpoIIIE protein for correct chromosome segregation. Molecular Microbiology 103, 504–517 (2017).

83. H. Veiga, A. Jousselin, S. Schäper, B. M. Saraiva, L. B. Marques, P. Reed, J. Wilton, P. M. Pereira, S. R. Filipe, M. G. Pinho, Cell division protein FtsK coordinates bacterial chromosome segregation and daughter cell separation in *Staphylococcus aureus*. The EMBO Journal 42, e112140 (2023).

84. C. Mercy, A. Ducret, J. Slager, J.-P. Lavergne, C. Freton, S. N. Nagarajan, P. S. Garcia, M.-F. Noirot-Gros, N. Dubarry, J. Nourikyan, J.-W. Veening, C. Grangeasse, RocS drives chromosome segregation and nucleoid protection in *Streptococcus pneumoniae*. Nature Microbiology 4, 1661–1670 (2019).

85. M. A. Schumacher, J. Lee, W. Zeng, Molecular insights into DNA binding and anchoring by the *Bacillus subtilis* sporulation kinetochore-like RacA protein. Nucleic Acids Research 44, 5438–5449 (2016).

86. S. Ben-Yehuda, M. Fujita, X. S. Liu, B. Gorbatyuk, D. Skoko, J. Yan, J. F. Marko, J. S. Liu, P. Eichenberger, D. Z. Rudner, R. Losick, Defining a centromere-like element in *Bacillus subtilis* by identifying the binding sites for the chromosome-anchoring protein RacA. Molecular Cell 17, 773–782 (2005).

87. S. Ben-Yehuda, D. Z. Rudner, R. Losick, RacA, a bacterial protein that anchors chromosomes to the cell poles. Science 299, 532–536 (2003).

88. A. N. Keller, Y. Xin, S. Boer, J. Reinhardt, R. Baker, L. K. Arciszewska, P. J. Lewis, D. J. Sherratt, J. Löwe, I. Grainge, Activation of Xer-recombination at dif: structural basis of the FtsKγ–XerD interaction. Scientific Reports 6, 33357 (2016).

89. J. Prikryl, E. C. Hendricks, P. L. Kuempel, DNA degradation in the terminus region of resolvase mutants of *Escherichia coli*, and suppression of this degradation and the Dif phenotype by *recD*. Biochimie 83, 171–176 (2001).

90. A. K. Sinha, A. Durand, J.-M. Desfontaines, I. Iurchenko, H. Auger, D. R. F. Leach, F.-X. Barre, B. Michel, Division-induced DNA double strand breaks in the chromosome terminus region of *Escherichia coli* lacking RecBCD DNA repair enzyme. PLOS Genetics 13, e1006895 (2017).

91. H. Veiga, M. G. Pinho, Inactivation of the SauI type I restriction-modification system is not sufficient to generate *Staphylococcus aureus* strains capable of efficiently accepting foreign DNA. Applied and Environmental Microbiology 75, 3034–3038 (2009).

92. T. Oshida, A. Tomasz, Isolation and characterization of a Tn551-autolysis mutant of *Staphylococcus aureus*. Journal of Bacteriology 174, 4952–4959 (1992).

93. M. Arnaud, A. Chastanet, M. Débarbouillé, New vector for efficient allelic replacement in naturally nontransformable, low-GC-content, gram-positive bacteria. Applied and Environmental Microbiology 70, 6887–6891 (2004).

94. J. Schindelin, I. Arganda-Carreras, E. Frise, V. Kaynig, M. Longair, T. Pietzsch, S. Preibisch, C. Rueden, S. Saalfeld, B. Schmid, J.-Y. Tinevez, D. J. White, V. Hartenstein, K. Eliceiri, P. Tomancak, A. Cardona, Fiji: an open-source platform for biological-image analysis. Nature Methods 9, 676–682 (2012).

95. R. F. Laine, K. L. Tosheva, N. Gustafsson, R. D. M. Gray, P. Almada, D. Albrecht, G. T. Risa, F. Hurtig, A.-C. Lindås, B. Baum, J. Mercer, C. Leterrier, P. M. Pereira, S. Culley, R. Henriques, NanoJ: a high-performance open-source super-resolution microscopy toolbox. Journal of Physics D 52, 163001 (2019).

96. U. Schmidt, M. Weigert, C. Broaddus, G. Myers, “Cell detection with star-convex polygons” in Medical Image Computing and Computer Assisted Intervention – MICCAI 2018, A. F. Frangi, J. A. Schnabel, C. Davatzikos, C. Alberola-López, G. Fichtinger, Eds. (Springer International Publishing, Cham, 2018), pp. 265–273.

97. J. D. Hunter, Matplotlib: a 2D graphics environment. Computing in Science & Engineering 9, 90–95 (2007).

98. Q. Liao, X. Wang, Using chromosome conformation capture combined with deep sequencing (Hi-C) to study genome organization in bacteria. Methods in Molecular Biology 2866, 231–243 (2025).

99. X. Wang, T. B. K. Le, B. R. Lajoie, J. Dekker, M. T. Laub, D. Z. Rudner, Condensin promotes the juxtaposition of DNA flanking its loading site in *Bacillus subtilis*. Genes & Development 29, 1661–1675 (2015).

100. I. R. Monk, I. M. Shah, M. Xu, M.-W. Tan, T. J. Foster, Transforming the untransformable: application of direct transformation to manipulate genetically *Staphylococcus aureus* and *Staphylococcus epidermidis*. mBio 3, e00277–11 (2012).

101. D. Nair, G. Memmi, D. Hernandez, J. Bard, M. Beaume, S. Gill, P. Francois, A. L. Cheung, Whole-genome sequencing of *Staphylococcus aureus* strain RN4220, a key laboratory strain used in virulence research, identifies mutations that affect not only virulence factors but also the fitness of the strain. Journal of Bacteriology 193, 2332–2335 (2011).

102. P. M. Pereira, H. Veiga, A. M. Jorge, M. G. Pinho, Fluorescent reporters for studies of cellular localization of proteins in *Staphylococcus aureus*. Applied and Environmental Microbiology 76, 4346–4353 (2010).

103. V. Vagner, E. Dervyn, S. D. Ehrlich, A vector for systematic gene inactivation in *Bacillus subtilis*. Microbiology 144 (Pt 11), 3097–3104 (1998).

